# An mTOR/RNA pol I axis shapes chromatin architecture in response to fasting

**DOI:** 10.1101/2023.07.22.550032

**Authors:** Nada Al-Refaie, Francesco Padovani, Francesca Binando, Johanna Hornung, Qiuxia Zhao, Benjamin D. Towbin, Elif Sarinay Cenik, Nicholas Stroustrup, Kurt M. Schmoller, Daphne S. Cabianca

**Affiliations:** Institute of Functional Epigenetics, Helmholtz Zentrum München, Neuherberg, Germany; Faculty of Medicine, Ludwig-Maximilians Universität München, Munich, Germany; Institute of Medical Sciences, University of Aberdeen, Aberdeen, UK; Department of Molecular Biosciences, University of Texas Austin, Austin, TX, USA; University of Bern, Bern, Switzerland; Centre for Genomic Regulation (CRG), The Barcelona Institute of Science and Technology, Barcelona, Spain; Universitat Pompeu Fabra (UPF), Barcelona, Spain

## Abstract

Chromatin architecture is a fundamental mediator of genome function. Fasting is a major environmental cue across the animal kingdom. Yet, how it impacts on 3D genome organization is unknown. Here, we show that fasting induces a reversible and large-scale spatial reorganization of chromatin in *C. elegans*. This fasting-induced 3D genome reorganization requires inhibition of the nutrient-sensing mTOR pathway, a major regulator of ribosome biogenesis. Remarkably, loss of transcription by RNA Pol I, but not RNA Pol II nor Pol III, induces a similar 3D genome reorganization in fed animals, and prevents the restoration of the fed-state architecture upon restoring nutrients to fasted animals.

Our work documents the first large-scale chromatin reorganization triggered by fasting and reveals that mTOR and RNA Pol I shape genome architecture in response to nutrients.

## Main Text

Regulation of chromatin architecture at multiple scales, from local DNA looping to larger-scale genome distribution at defined nuclear subcompartments, is a fundamental modulator of genome function (*1, 2*). During cell differentiation, 3D genome organization is modified in response to developmental cues (*3*). However, it remains largely unexplored whether, and how, environmental signals affect 3D chromatin organization.

Nutrients are a major environmental factor, since virtually all animals, including humans, are exposed to different diets and to feeding/fasting alternations. Perturbations of intracellular metabolism have been shown to regulate chromatin marks through changes in metabolites level, potentially influencing gene expression and organismal health (*4*). However, whether nutrients availability affects chromatin architecture at a larger scale in a multicellular organism is unknown. Here, we use *C. elegans* to investigate the impact of fasting on the large-scale genome organization using live imaging within tissues of intact animals, at the single-cell resolution.

### Fasting induces a tissue-specific, large-scale 3D reorganization of chromatin in the intestine of *C. elegans*

To investigate whether a complete lack of nutrients affects chromatin architecture, we exposed *C. elegans* larvae at the first larval stage (L1) to 12 hours of fasting. As a proxy for 3D genome organization, we monitored global distribution of five different fluorescently labeled histones: all H3 protein variants present in *C. elegans* HIS-72/H3.3, HIS-71/H3.3, HIS-6/H3.2 (*5, 6*), as well as HIS-24/H1.1 and HIS-1/H4. Live confocal imaging of fed and fasted worms showed that histones underwent a drastic spatial reorganization forming two “concentric rings” in all intestinal cell nuclei during fasting (Fig. 1A). To quantify chromatin distribution, we used the brightest HIS-72/H3.3 signal and segmented nuclei in 3D. For each nucleus, we averaged the fluorescence intensity profiles along the normalized center of nucleolar mass, which is known to be centrally positioned in intestinal nuclei (*7*), across all possible angles (fig. S1A, B). Consistent with an isotropic reorganization, we found nearly identical results when we compared this 3D analysis with one using only the xy plane, with the radial chromatin intensity shifting from a single peak in fed larvae, to two peaks during fasting (fig. S1A, B), allowing us to use 2D quantifications henceforth.

**Fig. 1.**
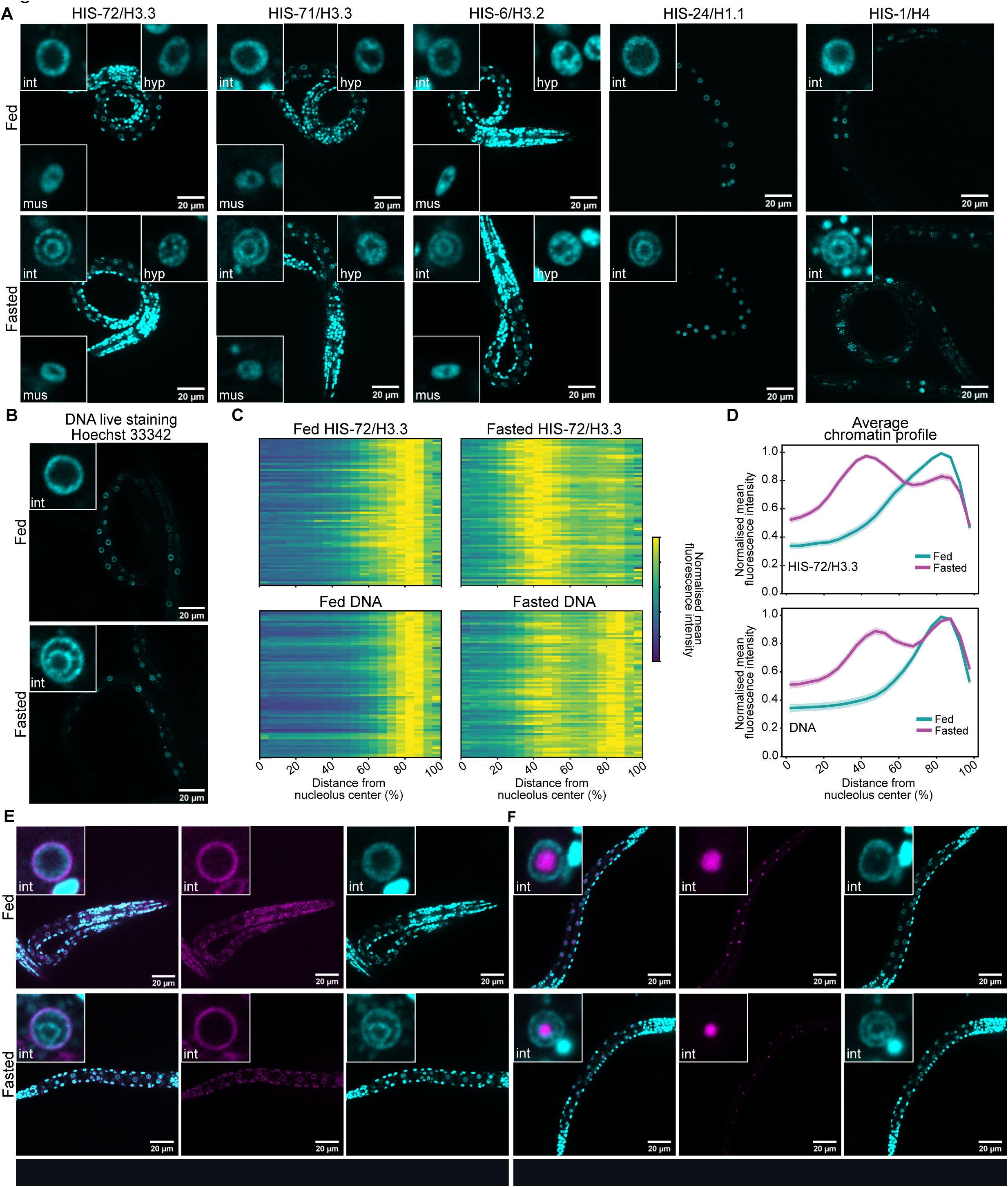
Fasting induces a large-scale genome reorganization in the *C. elegans* intestine. (**A**) Single focal planes of representative wt L1 larvae expressing the indicated fluorescently tagged histone, regularly fed (up) or fasted (bottom). Insets: zoom of single nucleus of the indicated tissue. hyp: hypodermis, int: intestine, mus: muscle. (**B**) as in (A) but live-stained with Hoechst 33342 to visualize DNA. (**C**) Heatmaps showing the radial fluorescence intensity profiles of HIS-72/H3.3-GFP (top) or Hoechst 33342-stained DNA (bottom) in intestinal nuclei of fed and fasted wt animals as a function of the relative distance from the nucleolus center, which occupies a central position (*7*). Each row corresponds to a single nucleus, segmented in 2D at its central plane. Radial fluorescence intensity profiles were averaged over all angles. 72 intestinal nuclei were analyzed in animals from 3 independent biological replicas. (**D**) Line plots of the averaged single nuclei profile shown in (C). The shaded area represents the 95% confidence interval of the mean profile. (**E**) Single focal planes of representative wt L1 larvae bearing HIS-72/ H3.3-GFP and EMR-1-mCherry in fed and fasted conditions. (**F**) as in (E), but expressing FIB-1-mCherry. Insets: zoom of single intestinal nucleus.

The fact that all histones tested reorganized similarly into two rings in intestinal cells, suggested that the genome at large, rather than specific subdomains, is spatially reorganized in response to fasting. Indeed, the same concentric rings are visible using the live imaging of DNA in fasted worms (Fig. 1B) and quantification of chromatin fluorescence distribution revealed comparable large-scale reorganizations in DNA- and HIS-72/H3.3-labeled nuclei (Fig. 1C, D).

We find that other tissues, such as hypodermis and muscle, did not reorganize the genome into two rings (Fig. 1A and fig. S1C-F), indicating that the 3D chromatin response is tissue-specific. To confirm that this spatial reorganization of chromatin is determined by the nutritional status of the organism, we showed that similar chromatin reorganizations occurred when animals were fed and fasted on plates or in liquid (Fig. 1A-D and fig. S1G-I), and over a wide range of physiological osmolarities (fig. S1J-L). Moreover, we did not observe a fasting-like 3D genome organization in heat- or cold-shock conditions (fig. S1M-O), indicating that the observed spatial reorganization of chromatin into two rings is not a general response to stress. Remarkably, adult worms in which intestinal cells no longer undergo DNA replication (*7*) also reorganized their genomes into concentric rings upon fasting (fig. S1P-R). This result argues that the newly identified 3D reorganization of the genome can occur in absence of cell division and is not restricted to specific larval stages.

The striking reorganization of the genome in the intestinal nuclei during fasting led us to hypothesize that these concentric rings might reflect an accumulation of chromatin at specific nuclear structures. Indeed, by conducting in vivo imaging, we found that the outer ring aligns with the nuclear envelope, as revealed by monitoring both chromatin and the nuclear envelope protein EMR-1/Emerin (Fig. 1E), while the inner ring encircled the single, centrally located nucleolus of intestinal cells (*7*), visualized by labeling its core component FIB-1/Fibrillarin (Fig. 1F). We note that chromatin was not enriched in direct proximity of the nucleolus in fed intestinal cells, in agreement with recent live imaging results in flies and human lymphocytes (*8*). Indeed, by measuring the radial chromatin intensity from the nucleolus towards the nuclear periphery in the fed state, we find that the signal is low at the edge of the nucleolus and increases moving towards the nuclear edge. In contrast, in fasted cells the chromatin signal is highest near the nucleolus (fig. S1S), confirming that the inner chromatin ring locates around the nucleolus.

### The fasting-induced chromatin rings are repressive compartments

The 3D organization of chromatin reflects the functional compartmentalization of the genome (*9*). Thus, we asked whether positioning of a gene at either the outer or the inner chromatin ring influences its expression. To this end, gene localization and expression need to be determined within single nuclei. Because standard fixation procedures disrupt the fasting-induced organization of chromatin (fig. S2A), as shown for other nuclear substructures (*10*), we chose to monitor a transcriptionally repressed (heterochromatic) and an active (euchromatic) reporter using a LacO/LacI-GFP-based strategy in living cells.

The cells of the *C. elegans* intestine can be divided into three subgroups that show distinct patterns of spatial gene positioning (*11*) and gene expression (*7, 12*). We focused on the most abundant of these three intestinal subgroups, namely the cells that occupy the middle section of the digestive tract (fig. S2B). To quantify positioning of the heterochromatic allele (fig. S2C), we used a zoning assay (*13*) (and fig. S2D) that scores the frequency with which a fluorescently tagged allele is found in one of three concentric zones of equal surface. As previously reported (*11*), the repressed, heterochromatin reporter is strongly enriched at the nuclear periphery in fed intestinal cells. We find that its distribution is not altered during fasting (fig. S2E).

Next, we quantified the position of a euchromatic gene (Fig. 2A), which was previously reported to be located at the nuclear interior in intestine (*14*), with respect not only to the nuclear periphery but also to the single, centrally located nucleolus. We found that during fasting the euchromatic allele shifted towards the nucleolus (Fig. 2B, C), consistent with a shift to the inner chromatin ring (Fig. 1F and fig. S1S). To determine the position and activity of a euchromatic reporter during fasting conditions, we used a trascriptionally active locus carrying the same promoter (*pha-4*) as before (Figure 2A), driving the expression of a histone-mCherry cassette, to allow the quantitation of expression while simultaneously visualizing the locus’ spatial distribution (Fig. 2D). As expected, in fed animals the active reporter predominantly occupied an internal position (Fig. 2E, F). During fasting, 24.1% (+/- 2.5%) of alleles are localized to the inner ring, and 26% (+/- 3.3%) are found at the outer ring (Fig. 2E, F), which is similar to the percentage with peripheral localization in fed animals. The majority of the active alleles, however, occupied the space between the rings (49.9% (+/- 4.7%)). To reveal whether allele positioning is associated with differences in expression, we defined the location of both alleles present in the diploid intestinal cells and quantified mCherry levels at the single nucleus level in fasted animals. Notably, the highest mCherry signal is measured in intestinal nuclei where both alleles are located between the two chromatin rings (Fig. 2G). The presence of one of the two alleles at either the inner or outer ring is associated with a diminished mCherry expression, and expression is lowest in nuclei in which both alleles are found at either one or the other chromatin ring (Fig. 2G). While we cannot exclude that allele position during fasting is pre-determined by chromatin features and/or expression levels that were established in the cells of fed larvae, these results indicate that the perinuclear and perinucleolar chromatin rings are repressive compartments.

**Fig. 2.**
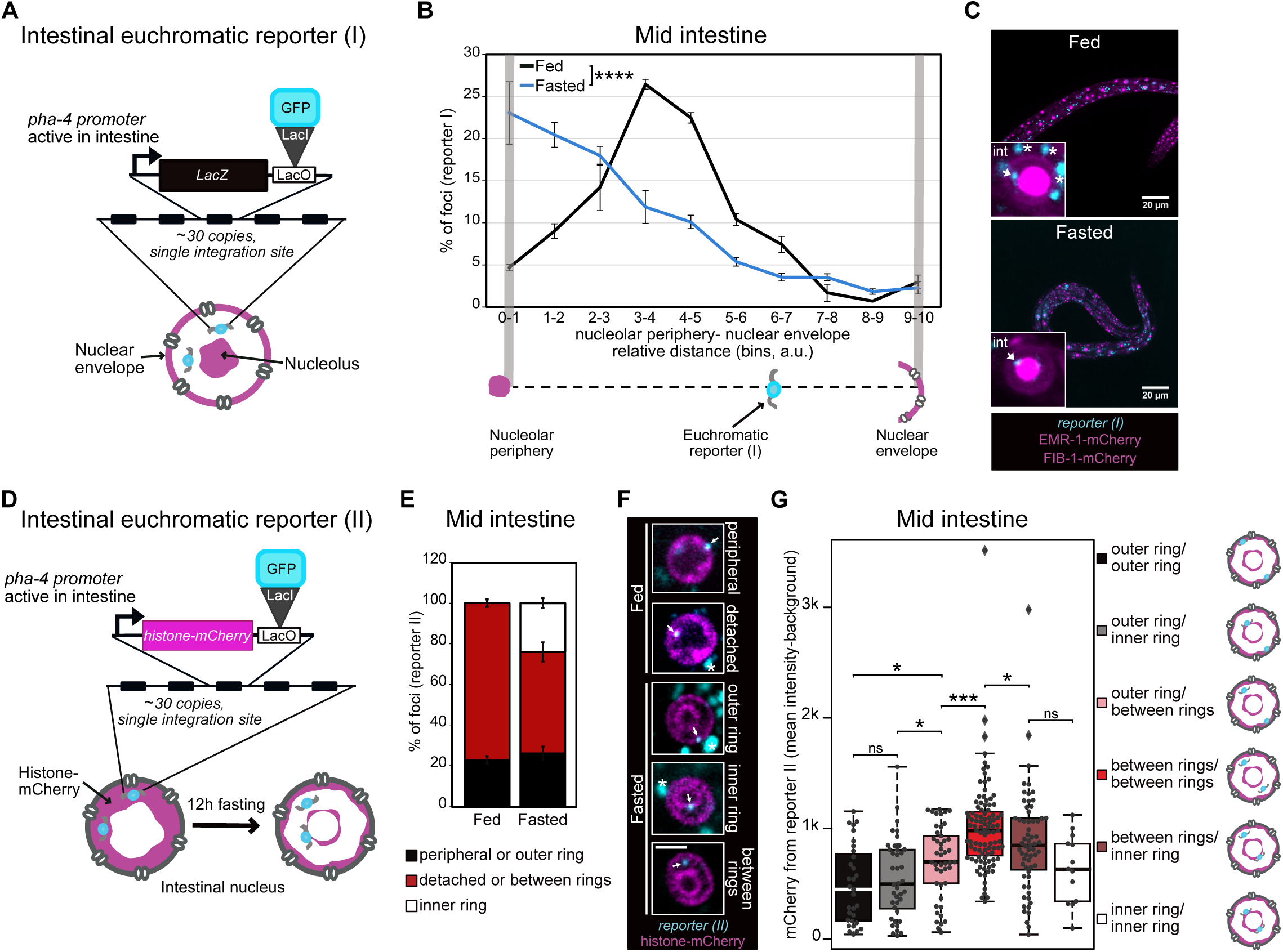
The inner and outer chromatin rings are repressive compartments. (**A**) Schematic representation of the euchromatin reporter (I) used in (B) and (C). (**B**) Line plots of the percentage of alleles represented in (A) found at the indicated relative position between the nucleolus and the nuclear envelope in fed and fasted animals. The average of 3 independent biological replicas are shown. Error bars are SEM. p value by χ^2^ is shown; **** indicates a p value <0.0001. See n and p value in table S2. (**C**) Single focal planes of representative wt L1 larvae expressing the indicated fluorescently-tagged markers, fed (top) or fasted (bottom). Insets: zoom of single intestinal nucleus. * mark autofluorescence from the intestinal cytoplasm. (**D**) Schematic representation of the euchromatin reporter (II) used in (E) (F) and (G). (**E**) Quantification of the sub-nuclear distribution of the active allele represented in (D) in the indicated locations in fed and fasted animals. The average of 5 independent biological replicas are shown. Error bars are SEM. See n values in table S2. (**F**) Single focal planes of representative single wt intestinal nuclei in L1 larvae expressing the indicated fluorescently-tagged markers, fed (top) or fasted (bottom) and with the indicated sub-nuclear location of the allele in (D). Scale bar indicates 2.5 µm, * marks autofluorescence from the intestinal cytoplasm. (**G**) Boxplots comparing the intensity of mCherry in single nuclei grouped based on the indicated localization of the two alleles, represented in (D). The results from 5 independent biological replicas are shown. Probability values from Wilcoxon rank sum tests to compare the indicated pairs are shown: p values < 0.05 and < 0.001 are indicated by * and ***, respectively. ns = not significant. See p values and n in table S2.

### The restoration of a fed-like genome architecture upon refeeding requires transcription by RNA Pol I

Because nutrient deprivation induces a reorganization of the 3D genome, we asked whether refeeding would reverse it. We therefore fasted L1 larvae for 12 hours, put them back on food and monitored the intestinal chromatin distribution over time. While after 5 minutes on food the genome is still organized into concentric rings (Fig. 3A, C, D), at later time points this configuration is lost. Indeed, refeeding for 30 minutes was sufficient to restore the normal “fed” 3D genome configuration (Fig. 3A, C, D). This argues that the spatial architecture of the genome of intestinal cells is dynamic and reflects the nutritional status of the organism.

**Fig. 3.**
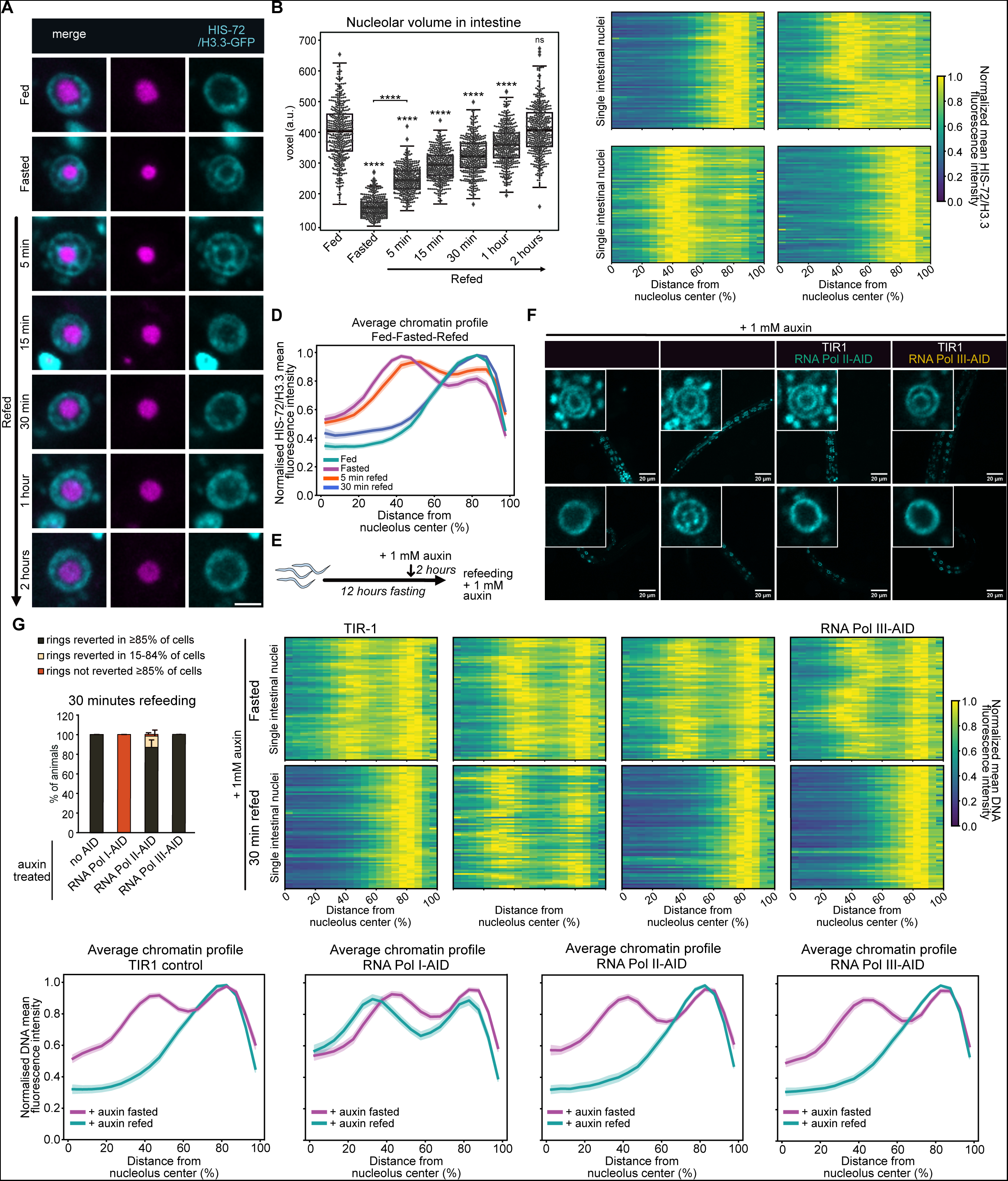
The restoration of a fed-like genome architecture upon refeeding requires RNA pol I transcription. (**A**) Single focal planes of representative single wt intestinal nuclei expressing HIS-72/H3.3-GFP and FIB-1-mCherry in L1 larvae that were fed, fasted or refed for the indicated time. Scale bar indicates 2.5 µm. (**B**) Boxplots comparing the volume of the nucleolus, measured with FIB-1-mCherry, in the intestine of larvae that were fed, fasted, or refed for the indicated time. Probability values from Wilcoxon rank sum tests comparing to fed are shown: **** indicates p value <0.0001. ns = not significant. See p values and n in table S2. (**C**) Heatmaps showing the radial fluorescence intensity profiles of HIS-72/H3.3-GFP in intestinal nuclei of wt animals of the indicated nutritional status as a function of the relative distance from the nucleolus center. Each row corresponds to a single nucleus, segmented in 2D at its central plane. Radial fluorescence intensity profiles were averaged over all angles. 72 intestinal nuclei were analyzed in animals from 3 independent biological replicas. (**D**) Line plots of the averaged single nuclei profile shown in (C). The shaded area represents the 95% confidence interval of the mean profile. (**E**) Schematic representation of the experimental timeline performed in (F-I) to test the requirement of transcriptional activities in reverting the chromatin rings by refeeding. (**F**) Single focal planes of representative L1 larvae live-stained with Hoechst 33342 expressing a ubiquitous TIR1 and either RPOA-2-AID (RNA Pol I-AID), RPB-2-AID (RNA Pol II-AID), RPC-1-AID (RNA Pol III-AID), or no AID as control (TIR1) fasted (up) or refed for 30 minutes (bottom) in presence of auxin, as outlined in (E). Insets: zoom of single intestinal nucleus. (**G**) Quantification of the percentage of the indicated AID-tagged animals (Pol I-, Pol II-Pol III- and no AID, as control) within the indicated categories for 3D chromatin organization in intestine after 30 minutes refeeding. (**H**) Heatmaps showing the radial fluorescence intensity profiles of DNA in fasted (top) or 30 minutes refed (bottom) in intestinal nuclei of animals expressing the indicated AID tag in presence of auxin as in (E), as a function of the relative distance from the nucleolus center. Each row corresponds to a single nucleus, segmented in 2D at its central plane. Radial fluorescence intensity profiles were averaged over all angles. 69-72 intestinal nuclei were analyzed in animals from 3 independent biological replicas. For RNA Pol II-AID 30 minutes refed, animals in proportions to their relative abundance within the 3D chromatin organization categories as in (G) were analyzed. (**I**) Line plots of the averaged single nuclei profile shown in (H). The shaded area represents the 95% confidence interval of the mean profile.

Intriguingly, it has been shown in several species that the volume of the nucleolus is reduced in the absence of nutrients (*15, 16*). Consistently, by using live imaging of animals expressing FIB-1/Fibrillarin-mCherry, we found that fasting led to a strong reduction in nucleolar volume in intestinal cells (Fig. 3B). Moreover, we found that upon refeeding, the nucleoli enlarge. Whereas 2 hours are required for the nucleolus to regain the size of fed animals, refeeding for only 5 minutes was sufficient to detect an increase in nucleolar volume. This argues that nucleolar size is regulated very early in the response to refeeding, which ultimately leads to the dispersion of the chromatin rings.

Transcription is a major driver of genome organization (*17–21*), and nucleolar size typically reflects the production of rRNA, with bigger nucleoli correlating with higher levels of RNA Pol I transcription (*22, 23*). However, nucleolar structure and size have also been reported to be regulated by RNA Pol II (*24, 25*). Therefore, we tested whether transcription by RNA Pol I, Pol II or Pol III has a role in restoring the fed-like 3D chromatin organization upon refeeding. To this end, we introduced AID (Auxin-Inducible-Degron (*26*)) and a fluorescent tag at the endogenous loci of the genes *rpoa-2*, *rpb-2* and *rpc-1* (corresponding to *POLR1B*, *POLR2B* and *POLR3A* in humans), which encode core subunits of RNA Pol I, Pol II and Pol III, respectively. The addition of auxin should lead to their acute degradation, selectively inhibiting the transcriptional activity of the targeted RNA polymerase. One hour on auxin rendered the RNA Pol III- and II-specific core subunits RPC-1 and RPB-2 undetectable in intestine, while about four hours were needed for the RNA Pol I-specific component RPOA-2 (fig. S3).

We fasted larvae to induce the 3D genome reorganization into two rings, added auxin during the last two hours of nutrient deprivation (Fig. 3E) and subsequently, placed the worms on food- and auxin-containing plates for 30 minutes. The 3D chromatin architecture reverted to the fed-like state in control animals (TIR1 expressing only), as well as in those depleted for RNA Pol II and III subunits. However, the animals depleted for the RNA Pol I core subunit were unable to restore the fed-like chromatin configuration, retaining the concentric rings of fasted cells despite feeding (Fig. 3F-I). Because DNA staining can only be obtained if worms eat Hoechst 33342-containing bacteria, we can exclude that the retention of the chromatin rings in RNA Pol I-depleted animals stems from a failure to eat. From this, we conclude that RNA Pol I, but not Pol II nor Pol III, is necessary to re-establish the fed-like genome architecture upon refeeding.

### Inhibition of RNA Pol I transcription is sufficient to induce a fasting-like chromatin architecture in the intestine of fed animals

When nutrients are scarce, cells reduce ribosome synthesis by downregulating RNA Pol I transcription (*27*). A fasting-induced decrease in RNA Pol I activity is consistent with the reduced nucleolar volume that we observe in fasted intestinal nuclei (Fig. 3B) and the shrinkage of the nucleolus induced by RPOA-2 degradation (fig. S4A). However, RNA Pol I is only one of many targets affected by fasting (*28*). To determine if inhibition of RNA Pol I transcription per se can disrupt the normal 3D genome organization in fed animals, we monitored chromatin distribution in fed larvae upon degradation of RPOA-2. Strikingly, even though complete RPOA-2 degradation in fed worms required nearly 4 hours on auxin (fig. S3A, B), 1 hour on auxin was sufficient to induce strong changes in chromatin distribution (Fig 4A, B). After 3 hours on auxin, the concentric chromatin rings appeared in the majority of larvae (Fig. 4A, B) and a fasting-like radial spatial reorganization of chromatin could be observed when averaging over all cells at this time point (Fig. 4C, D). After 5 hours, 100% of these animals had converted the fed chromatin distribution into two concentric chromatin rings, in more than 80% of the intestinal cells (Fig. 4A, B). To confirm this result, we inhibited RNA Pol I transcription by the addition of Actinomycin D, and found that this also induces a fasting-like reorganization of chromatin in intestine of fed animals (fig. S4B-D).

**Fig. 4.**
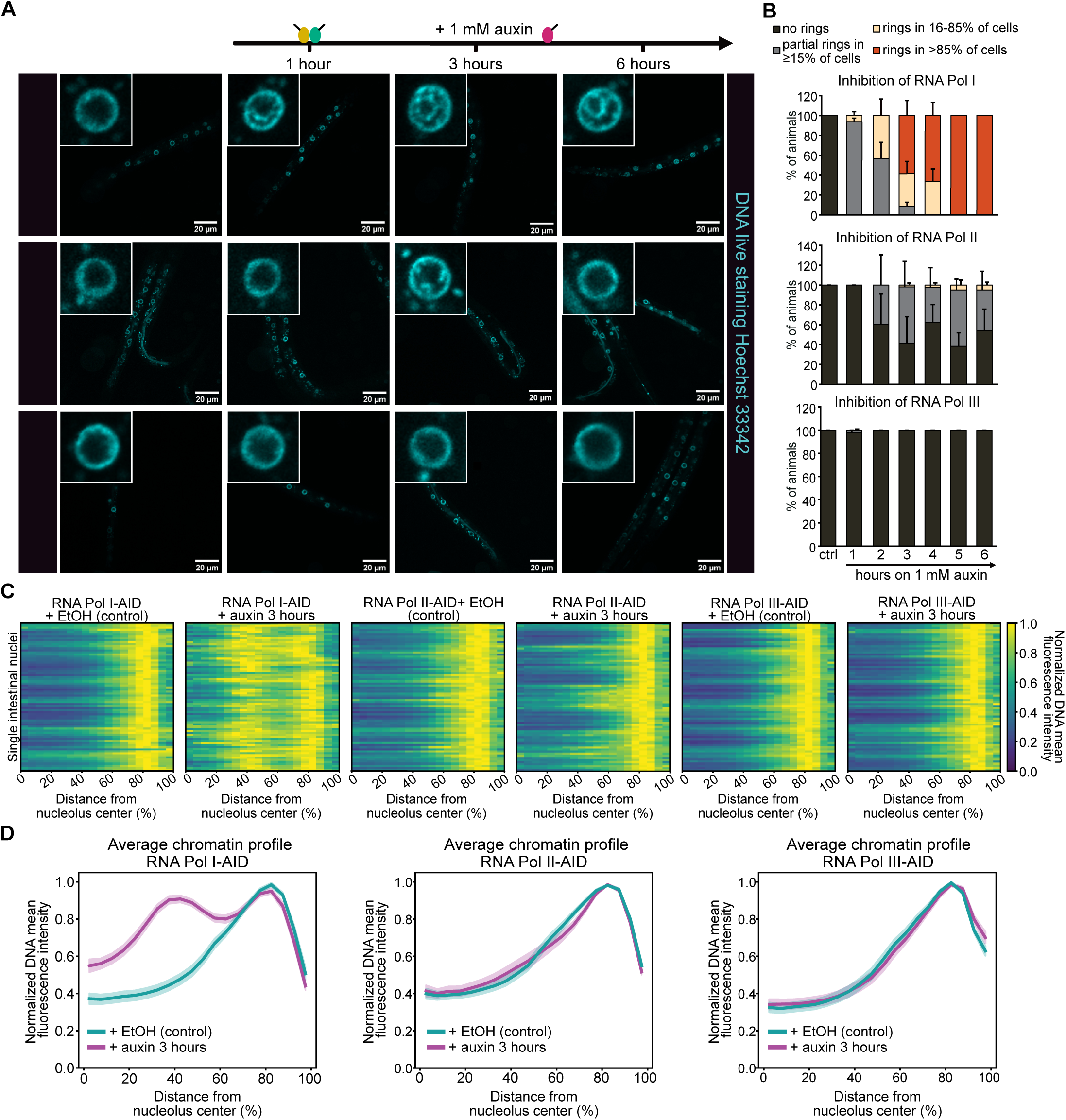
Inhibition of RNA Pol I activity is sufficient to induce a fasting-like chromatin architecture in the intestine of fed animals. (**A**) Top, schematic representation of the degradation kinetics of the core subunits RPC-1 (RNA Pol III), RPB-2 (RNA Pol II) and RPOA-2 (RNA Pol I) in intestine, as quantified in fig. S3. Bottom, single focal planes of representative fed L1 larvae live-stained with Hoechst 33342 to monitor DNA, ubiquitously expressing TIR-1 and the indicated core subunit endogenously-tagged with degron (AID) are shown during EtOH exposure, as control, and after 1, 3 and 6 hours of 1 mM auxin exposure. Insets: zoom of single nucleus of the intestine. (**B**) Quantification of the percentage of the indicated AID-tagged animals (RNA Pol I-, Pol II- and Pol III -) within the indicated categories for 3D chromatin organization in intestine, at the indicated time points of auxin exposure. (**C**) Heatmaps showing the radial fluorescence intensity profiles of DNA in intestinal nuclei of the indicated fed, degron-tagged animals in presence of EtOH, as control, or 3 hours on 1 mM auxin as a function of the relative distance from the nucleolus center. Each row corresponds to a single nucleus, segmented in 2D at its central plane. Radial fluorescence intensity profiles were averaged over all angles. 72 intestinal nuclei were analyzed in animals from 3 independent biological replicas. For RNA Pol I- and Pol II-AID at 3 hours of auxin exposure, animals in proportions to their relative abundance within the 3D chromatin organization categories as in (B) were analyzed. (**D**) Line plots of the averaged single nuclei profile shown in (C). The shaded area represents the 95% confidence interval of the mean profile.

To test whether this effect is specific to RNA Pol I transcription, we blocked the activity of RNA Pol II and III in fed animals, using the RBP-2- and RPC-1-degron strains, respectively. Remarkably, after 1 hour on auxin we detected no changes in the global distribution of chromatin in intestine (Fig. 4A, B), despite both core subunits being fully degraded at this time point (fig. S3C-F). While loss of RNA Pol III transcription did not affect chromatin organization even after 3 or 5 hours of auxin exposure (Fig 4A-D), variable changes in chromatin distribution could be scored upon RNA Pol II inhibition, with concentric rings of chromatin being detected in about 2-5% of animals, depending on the time point (Fig. 4B). Still, the average chromatin organization was largely unaffected by RNA Pol II inhibition (Fig. 4C, D). We conclude that transcription by RNA Pol I is necessary to maintain the chromatin architecture typical of the fed state in intestinal cells as its inhibition, and not that of RNA Pol II nor III, is sufficient to induce a fasted-like chromatin reorganization in the intestine of fed animals.

### mTORC1 signaling is necessary and sufficient to regulate 3D genome architecture in response to nutrients

The regulation of the large-scale chromatin organization in intestine by fasting, suggests that nutrient-sensing signaling pathways might be implicated. We investigated the role of monophosphate (AMP)-activated kinase (AMPK), which is activated by a reduction in the ATP/AMP ratio (*29*) under poor nutrient conditions. Despite causing a decreased survival to prolonged starvation (*30*), mutating the two *C. elegans* AMPK catalytic subunits *aak-1* and *aak-2* did not alter the 3D chromatin reorganization of intestinal cells during fasting (fig. S5A-C), suggesting that the AMPK signaling pathway is not involved.

The mTOR, or mechanistic target of rapamycin, signal-transduction pathway allows cells to adjust their protein biosynthetic capacity to nutrient availability (*27*). In particular, mTOR is active in rich nutrient conditions and inactive in absence of nutrients. RAPTOR, termed DAF-15 in worms (*31*), is a critical effector of mTORC1, the mTOR-containing complex that regulates protein synthesis and ribosome biogenesis in response to nutrient availability (*32*). We thus asked what is the effect of mTORC1 inhibition on the spatial organization of chromatin in fed animals. We found that depleting DAF-15 using the auxin-inducible-degradation system was sufficient to induce a partial reorganization of chromatin into concentric rings in the intestinal cells of fed animals (Fig. 5A-D). In mammals, the inactivation of mTORC1 in absence of nutrients is counteracted by a Q66L substitution in RagA, a protein of the Rag family of GTPases that acts upstream of mTORC1. This mutation impairs GTP hydrolysis, rendering RagA constitutively active, thus keeping mTOR signaling active even when nutrients are lacking (*33, 34*). In *C. elegans*, a similar phenotype is observed when RAGA-1, homolog of RagA, carries a Q63L mutation (*35*). We introduced the Q63L mutation in the endogenous *raga-1* gene, thus generating a constitutively active gain of function mutant, *raga-1^GF^*. These *raga-1^GF^* mutants had larger nucleoli in intestinal cells compared to wild-type (Fig. 5E), consistent with mTOR being an evolutionary conserved regulator of nucleolar size (*15, 16, 36*), potentially through the direct regulation of rDNA transcription (*37*). Importantly, keeping mTORC1 active during fasting antagonized the reorganization of chromatin into concentric rings in intestine (Fig. 5F-I), revealing that mTOR inactivation is required for the spatial reorganization of the genome induced by fasting. Remarkably, expressing RAGA-1GF does not antagonize the formation of the concentric rings upon RNA Pol I inhibition (Fig. 5J-L), showing that RNA Pol I transcription acts downstream of mTORC1 in the regulation of the 3D chromatin architecture by nutrients (Fig. 5M).

**Fig. 5.**
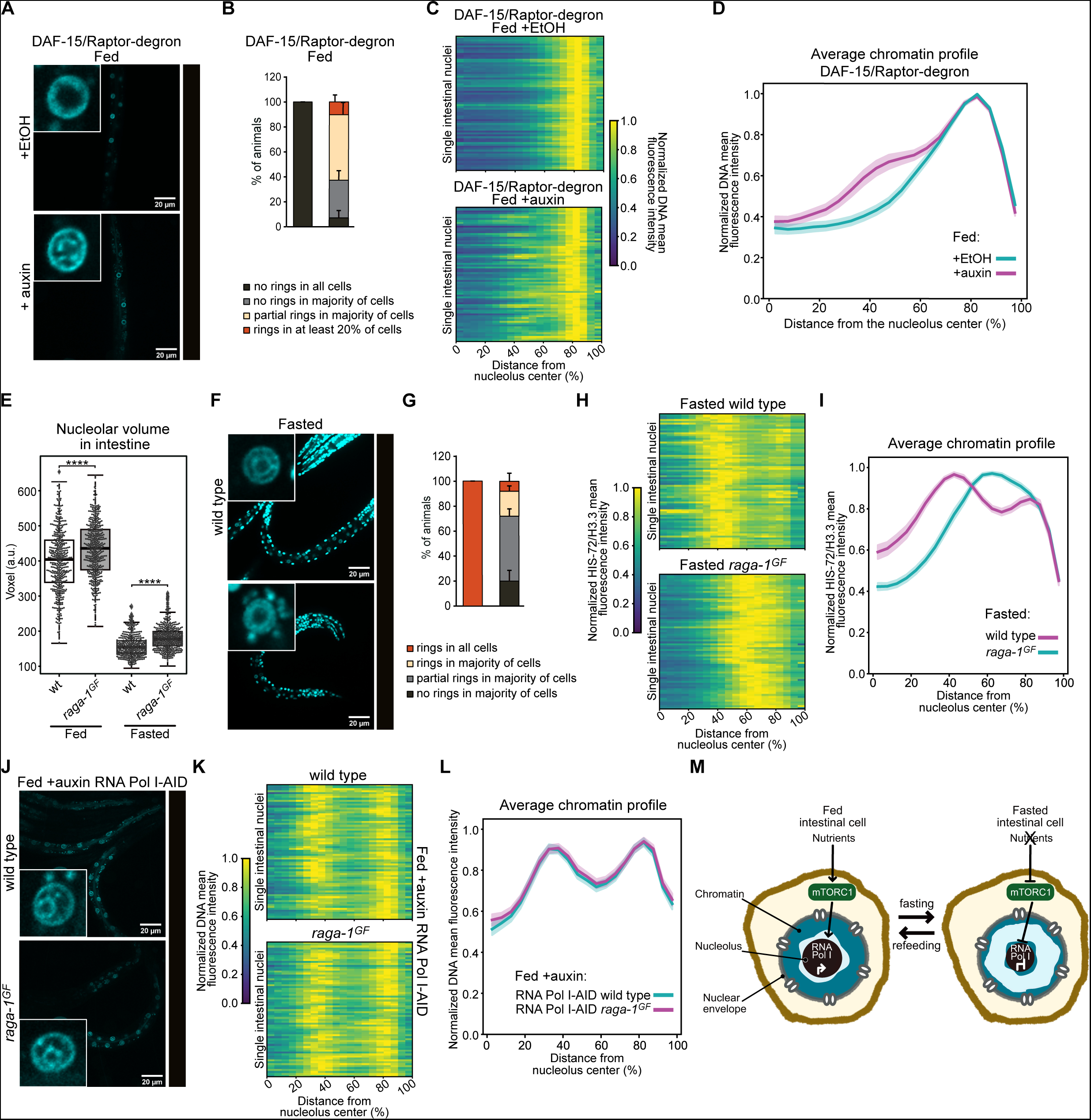
mTORC1 signaling is necessary and sufficient to regulate 3D genome architecture in response to nutrients in intestine. (**A**) Single focal planes of representative, fed L1 larvae expressing an endogenously degron-tagged DAF-15/Raptor treated with 1 mM auxin to induce its degradation, or EtOH, as control, and where the DNA has been stained with Hoechst 33342. Insets: zoom of single nucleus of the intestine. (**B**) Quantification of the percentage of fed DAF-15/Raptor-degron tagged animals treated with 1 mM auxin or EtOH within the indicated categories for 3D chromatin organization in intestine. (**C**) Heatmaps showing the radial fluorescence intensity profiles of DNA in intestinal nuclei of fed DAF-15/Raptor-degron animals treated with EtOH (top) or auxin (bottom) as a function of the relative distance from the nucleolus center. Each row corresponds to a single nucleus, segmented in 2D at its central plane. Radial fluorescence intensity profiles were averaged over all angles. 71-72 intestinal nuclei were analyzed in animals from 3 independent biological replicas. For auxin-treated individuals, animals in proportions to their relative abundance within the chromatin organization categories as in (B) were analyzed. (**D**) Line plots of the averaged single nuclei profile shown in (C). The shaded area represents the 95% confidence interval of the mean profile. (**E**) Boxplots comparing the volume of the nucleolus, measured with FIB-1-mCherry, in the intestine of fed and fasted wt and *raga-1^GF^* larvae. Probability values from Wilcoxon rank sum tests comparing to wt are shown: **** indicates p value < 0.0001, respectively. See p values and n in table S2. (**F**) as in (A) but showing fasted wt and *raga-1^GF^* animals expressing HIS-73/H3.3-GFP. Insets: zoom of single nucleus of the intestine. (**G**) Quantification of the percentage of fasted wt and *raga-1^GF^* animals in the indicated categories for 3D chromatin organization in intestine. (**H**) as in (C) but for fasted wt and *raga-1^GF^* animals. 72 intestinal nuclei were analyzed in animals from 3-4 independent biological replicas. For *raga-1^GF^*, larvae in proportions to their relative abundance within the chromatin organization categories as in (G) were analyzed. (**I**) Line plots as in (D) but for fasted wt and raga-1*^GF^* larvae. (**J**) as in (A) but showing fed wt and *raga-1^GF^* animals expressing RNA Pol I degron-tagged (RNA Pol I-AID) treated with 1 mM auxin for 5 hours and live-stained with Hoechst 33342 to monitor DNA. Insets: zoom of single nucleus of the intestine. (**K**) as in (C) but for fed wt (top) and *raga-1^GF^* (bottom) animals expressing RNA Pol I degron-tagged (RNA Pol I-AID) treated with 1 mM auxin for 5 hours. 70 intestinal nuclei were analyzed in animals from 3 independent biological replicas. (**L**) Line plots as in (D) but for fed wt and *raga-1^GF^* animals expressing RNA Pol I degron-tagged (RNA Pol I-AID) treated with 1mM auxin for 5 hours. (**M**) Model: in fed animals, mTORC1 promotes transcription by RNA Pol I within the nucleolus. During fasting, mTORC1 inactivation represses RNA Pol I transcription and reduces nucleolar size, thus promoting the reorganization of the genome which becomes enriched at the nuclear and nucleolar periphery.

## Discussion

Connections between cellular metabolism and chromatin architecture have been studied primarily in yeast (*38*) or in isolated cells or tissues (*39*). In this work, we describe for the first time, how the nutritional state alters the large-scale spatial organization of the genome of a living multicellular organism.

By using live confocal microscopy in *C. elegans*, we discovered that fasting triggers a reversible (Fig. 3A, C, D) and large-scale 3D genome reorganization specifically in intestine (Fig 1A and S1C-F). Based on our results, we propose that the 3D genome architecture of intestinal cells is regulated by mTOR signaling in response to nutrients through the modulation of RNA Pol I activity (model in Fig. 5M). We find that RNA Pol I acts downstream of mTOR (Fig. 5J-L) and has a critical and unique role in shaping the 3D genome architecture in intestine. In fact, its inhibition, but not that of Pol II or III, in fed animals is sufficient to mimic fasting and induce the two chromatin rings organization (Fig. 4). In addition, transcription by RNA Pol I, but not Pol II or Pol III, is essential to reverse the chromatin rings upon refeeding (Fig. 3F-I)

Our work suggests that the role of mTOR and RNA Pol I in regulating the 3D genome might differ depending on the cell type and might be determined by specific cellular properties. Besides its essential role in nutrient absorption and in the production of enzymes to be secreted into the gut lumen, the *C. elegans* intestine carries out functions that in mammals are executed by distinct organs, such as detoxification of chemicals and fat storage and, possibly due to their high level of protein synthesis, have large nucleoli compared to other *C. elegans* cell types (*40*). During fasting, the nucleolar volume in intestinal cells drastically decreases (Fig. 3A, B) altering the general architecture of the nucleus and potentially impacting on the 3D chromatin organization.

There is growing evidence that environmental cues in the form of diet and lifestyle contribute to the onset of metabolic diseases in humans via epigenetic mechanisms (*41*). While a role for chromatin architecture in this process remains undetermined, our work unveils that nutritional stimuli from the environment alter the spatial organization of the genome, adding an additional layer of complexity to the regulation of genome architecture, which may contribute to health and diseased states.

## Acknowledgements

We thank Robert Schneider, Maria-Elena Torres-Padilla and Susan M. Gasser for critical reading of the manuscript, the labs of Helge Grosshans and Florian Steiner, Ryan Gleason in Xin Cheńs lab, the Caenorhabditis Genetics Center (CGC), funded by NIH Office of Research Infrastructure Programs (grant no. P40 OD010440CGC), and the C. elegans Gene Knockout Project at the Oklahoma Medical Research Foundation for sharing strains. We thank SunyBiotech for support in generating the rpc-1-mNeonGreen-AID allele. D. S. C. thanks Helmholtz Munich for support.

## Funding

The German Research Foundation (Deutsche Forschungsgemeinschaft, DFG) SFB 1064, collaborative research center in chromatin dynamics (DSC)

The German Research Foundation (Deutsche Forschungsgemeinschaft, DFG) SPP 2202 Priority Programme “Spatial Genome Architecture in Development and Disease” (DSC)

National Institutes of Health grant 5R35GM138340-03 (ESC)

The Welch Foundation F-2133-20230405 (ESC)

Human Frontier Science Program (career development award) (KMS)

The Swiss National Science Foundation (SNSF) in the form of an Eccellenza Professorial Fellowship (PCEFP3_181204) (BDT)

European Research Council under the European Union’s Horizon 2020 Research and Innovation Programme (grant agreement no. 852201) (NS)

The Spanish Ministry of Economy, Industry and Competitiveness to the EMBL partnership, the Centro de Excelencia Severo Ochoa (CEX2020-001049-S, MCIN/AEI/10.13039/501100011033), the CERCA Programme/Generalitat de Catalunya (NS)

The Spanish Ministry of Economy, Industry and Competitiveness Excelencia award PID2020-115189GB-I00 (NS)

## Author contributions

Conceptualization: NAR, DSC

Methodology: FP, KMS, DSC

Software: FP

Investigation: NAR, JH, FB, DSC

Visualization: FP, NAR, DSC, KMS

Validation: NAR, JH, FB

Resources: QZ, ESC, BDT, NS

Funding acquisition: DSC, KMS, ESC, NS, BDT

Supervision: DSC

Writing – original draft: NAR, DSC

Writing – review & editing: DSC, NAR, KMS, BDT

## Data and materials availability

All data are available in the main text or the supplementary materials and are available upon request. All code is available on GitHub at the following links: https://github.com/SchmollerLab/SeelMito and https://github.com/ElpadoCan/ChromRings

Strains generated in this study are available upon reasonable request to the corresponding author. Strains GW429, GW1056 and strains carrying the *raga-1^GF^* allele may be subject to an MTA.

## Materials and Methods

### *C. elegans* maintenance and strains

Nematodes were grown with *Escherichia coli* OP50 bacteria on NGM agar plates at 20°C except where otherwise stated. All strains used in this study are listed in table S1.

### Constructs and strains

Endogenously tagged *rpc-1* at the C-terminus with STSGGSGGTGGS-mNeonGreen-GSAGSA-degron was obtained by CRISPR/Cas9 from the company SunyBiotech.

For the HIS-1-GFP fusion construct, the *his-1* gene was amplified from N2 worm genomic DNA and fused by PCR to GFP, which contained introns. *ges-1* promoter and *unc-54* 3’ UTR were amplified from N2 genomic DNA. The final plasmid construct was generated by MultiSite Gateway® cloning (Invitrogen). The strain expressing HIS-1/H4-GFP was made using the MosSCI technique (*42*). The transgene was inserted into ttTi5605 on Chr II.

The *degron-GFP* tagged *rpoa-2* allele was constructed as described (*43*) using Cas9 protein driven by *eft-3* promoter in pDD162 and gRNA targeting a genomic sequence in the N-terminus of *rpoa-2* in pRR13, a derivative of pRB1017, an empty vector for gRNA cloning. *degron-GFP-c1^sec^3xflag* repair template was constructed for generating the knock-in into the N terminus of the *rpoa-2* gene. The 5’ and 3’ homology arms 751 bp upstream of *rpoa-2* start codon and 566 bp downstream of start codon were used to replace the ccdB cassette in *degron-GFP-c1^sec^3xflag* repair template. Each knock-in was isolated via hygromycin selection and the SEC was then excised by heat-shock to produce *degron::GFP::rpoa-2* strain.

### Feeding, fasting and refeeding

Worms were maintained well-fed on OP50 at 20°C for at least two generations.

Fed L1s were obtained by washing plates of mixed stages animals twice with M9 buffer to remove adults and larvae, the washes were performed with swirling very gently to avoid removing the bacteria. To obtain synchronized L1s, the embryos that remained on the plate were allowed to hatch for 2 hours.

For L1s fasting, embryos were isolated by standard hypochlorite treatment and maintained in M9 buffer on a roller at 20°C. L1 larvae hatch approximately 12 hours after hypochlorite treatment (*44* and our own observation), and hence, this time point was considered 0 hours of fasting. Consequently, 12 hours of fasting corresponds to 24 hours after hypochlorite treatment.

For fasting of adults, worms were synchronized at the L1 stage by standard hypochlorite treatment and grown on OP50-seeded plates until day 1 of adulthood. Next, adults were collected by washing off the plates with M9 buffer. A fraction of worms was immediately prepared for imaging of the fed state, the rest were washed 3 times with M9 buffer for 10 minutes to remove remaining bacteria and fasted in M9 buffer for 12 hours on a roller at 20°C.

Refeeding was performed by adding 12-hours fasted L1 on NGM plates seeded with OP50 for the indicated time. Then, worms were immediately collected and prepared for imaging.

For feeding in liquid, in order to obtain fed and synchronized L1 larvae that were maintained in liquid for the same duration as the fasted animals, embryos obtained from hypochlorite treatment were kept in M9 buffer for 21 hours. Next, L1s were pelleted and M9 buffer was removed and S-basal complete medium supplemented with 6 mg/ml *E.coli* OP50 was added. The larvae were subsequently fed in the liquid culture for 3 hours before imaging, reaching a total of 24 hours in liquid.

For fasting on plate, isolated embryos obtained by hypochlorite treatment were placed onto M9 agarose plates without any bacteria for 24 hours, so that L1s were fasted for an average of 12 hours as described above, for fasting in liquid.

### DNA staining

To prepare the staining solution, Hoechst 33342 (ChemCruz, 20 mM stock) was diluted 1:1000 in M9 buffer, resulting in a final concentration of 20 µM. Next, the staining solution was added on OP50 bacteria, previously seeded and dried on NGM plates, making sure that all bacteria are covered. Next, the plates were left to dry in the dark.

To perform DNA live staining in fed larvae, L1s were obtained by washing plates of mixed stages and waiting 2 hours for embryos to hatch. Hatched L1s were collected and added to these plates directly on the food and allowed to feed for 3 hours on Hoechst 33342-stained bacteria.

Staining of the intestinal DNA in fasted worms was achieved by first feeding L1s with Hoechst 33342-stained bacteria for 3 hours, as described above. Next, L1s were collected using M9, washed three times in M9 for 10 minutes to remove the bacteria, and subsequently fasted in M9 buffer for 6 hours.

### Auxin stock and plates

Auxin 3-Indoleacetic acid (IAA) (Sigma-Aldrich) was dissolved in ethanol to prepare a 57 mM stock solution and stored at 4 °C. Auxin was added to NGM plates to obtain a final concentration of 1 mM. Control plates contained an equivalent amount of ethanol (1.75 % final concentration). Both auxin and control plates were seeded with 250 µl of OP50 culture grown overnight in LB.

### Auxin treatment

#### Degradation of polymerase’ subunits

Degradation of polymerasé subunits in strains expressing RPB-2-AID, RPC-1-AID and RPOA-2-AID was obtained as follows: plates containing a mixture of worm stages were washed twice with M9 buffer to eliminate all stages except embryos, which adhere to the plates. The washes were performed by swirling delicately to retain a layer of OP50 intact. Next, freshly hatched L1 s were collected after 2 hours and added onto auxin or ethanol plates containing Hoechst 33342-stained OP50 bacteria. L1s were maintained on auxin plates for the indicated amount of time, and on ethanol plates for 3 hours, before imaging.

For the refeeding experiments in absence of the polymerases core subunits, embryos were isolated using hypochlorite treatment and left in M9 buffer to allow L1s to hatch in absence of food. After a fasting period of 10 hours (22 hours post-bleaching), an aliquot of L1s was DNA live-stained to verify the formation of the chromatin rings in intestine, by microscopy. Briefly, an aliquot of 10 hours-fasted L1s was incubated in M9 supplemented with 1 mM auxin and 40 µM Hoechst 33342 for 2 hours in a 15 ml tube covered with aluminum foil to avoid light exposure, next, L1s were imaged.

In parallel, auxin was added to the remaining 10 hours-fasted L1s at a final concentration of 1 mM in M9 buffer. Next, 2 hours after the addition of auxin (12 hours of fasting), L1s were transferred to auxin plates seeded with OP50 bacteria stained with Hoechst 33342. Next, L1s were fed for 30 minutes and imaged.

#### Degradation of DAF-15

Embryos isolated by hypochlorite treatment were washed 3 times in M9 and directly plated onto auxin or ethanol plates, seeded with OP50 bacteria. After 21 hours, hatched L1s were moved onto auxin or ethanol plates, respectively, seeded with Hoechst 33342-stained OP50. Since the absorption of auxin is less efficient in embryos due to their eggshell (*45*) and because L1s hatch approximately 12 hours after hypochlorite treatment of embryos (Webster et al., 2022), 24 hours of auxin treatment started as embryos was considered to be equivalent to 12 hours of auxin exposure in L1s. Degradation of DAF-15-AID upon auxin treatment was previously performed and found to require 1 hour on 1 mM auxin (*46*).

### Microscopy

Microscopy was carried out using a live-cell imaging system (Confocal Spinning Disk Microscope) from Visitron Systems GmbH, equipped as follows: Nikon Eclipse Ti2 microscope + Plan Apo λ 100X/1.45 oil objective, Plan apo λ 60X/1.40 oil objective, Yokogawa CSU-W1 confocal scanner unit, VS-Homogenizer, EMCCD camera [Andor - iXon Series], and VisiView software for acquisition.

Live microscopy was carried out on 2% agarose pads supplemented with 0.15 % sodium azide (Interchim) to paralyze the worms, as previously described (*14*). All Images were acquired with the Plan Apo λ 100X/1.45 oil objective except for adults which were acquired with the Plan apo λ 60X/1.40 oil immersion objective. For each image, a range of stacks, from 50 to 120 stacks depending on the stage, were captured with a z-spacing of 200 nm.

In fig. S2A left, fasted L1s were fixed with 4% paraformaldehyde in PBS for 5 minutes and washed with PBS containing 30 mM glycine (PBS-G) for 10 minutes at room temperature. Next, worms were permeabilized with acetone at -20°C for 1 minute, and subsequently washed with PBS-G. For staining, a solution was prepared by adding 2 drops of EasyProbe-Hoechst 33342 Live Cell Stain (GeneCopoeia) into 1 ml of M9 buffer. Next, the fixed worms were incubated in this staining solution for 15 minutes. After staining, a final 5 minutes wash with PBS-G was performed before preparing the samples for imaging on an agarose pad as described above but without sodium azide. In fig. S2A right, fasted L1s were spotted on poly-L-lysine–covered slides and allowed to settle for few minutes. Next, coverslips were applied and slides were snap-frozen on dry ice for 25 minutes. To permeabilize worms, samples were then freeze-cracked by flicking the coverslips and immersed immediately in 100% ice-cold methanol for 10 seconds. Next, the slides were transferred to a fixing solution (0.08 M Hepes pH 6.9, 1.6 mM MgSO4, 0.8 mM EGTA, 3.7% formaldehyde, in PBS) for 10 minutes. After fixation, slides were washed three times with TBS-T (TBS with 0.1% Tween-20). To visualize DNA, samples were incubated with 10 µM Hoechst 33342 (ChemCruz) for 2 hours, washed once with TBS-T, mounted with 80% glycerol in PBS and imaged.

### Image analyses

#### Fluorescence intensities

Fluorescence intensities of GFP, mNeonGreen and mCherry were measured using Fiji/ImageJ (*47, 48*). For the polymeraseśs subunits degradation experiments shown in figs. S3 and S4A, for each strain, images of the different treatments were acquired with the same settings. Intestinal nuclei were selected as region of interest based on Hoechst 33342 signal. Within each nucleus, quantitation of GFP or mNeonGreen mean signal intensity on focal stack images was done selecting the brightest plane and subtracting the average background of the corresponding image. In Fig. 2G, in mid intestinal nuclei, quantitation of the mCherry mean signal intensity on focal stack images was done selecting the central plane and subtracting the average background of the corresponding image.

#### Chromatin profiling

Intestinal nuclei were manually segmented in 2D or in 3D using Cell-ACDC (*49*) based on HIS-72-GFP or Hoechst 33342 signal. To extract the intensity profiles shown in Figs 1D, 3D, 3I, 4D, 5D, 5I, 5L, S1B, S1E, S1F, S1I, S1L, S1O, S1R, S4D, S5C, we implemented a custom routine written in Python. The analysis steps are the following: 1) Manual annotation of the center of the inner dark area of the nuclei using Cell-ACDC. This is assumed to be the center of the nucleolus. 2) Determination of the nuclei contours from the segmented objects (using the function from OpenCV package called ‘findContours’). 3) Extraction of the intensity profiles from the center determined in the previous step and all the points on the contour. 4) Normalization of each profile using the distance from the center to the contour point (0% center, 100% contour). 5) Binning the intensities into 5% width bins (e.g., intensities at any distance in the ranges 0-5%, 5-10%, 10-15%, etc. were considered being at 2.5%, 7.5%, 12.5% (bin center) etc. distance from the center). 6) Averaging of the binned normalized profiles along the distance to obtain the single-nuclei average profiles shown as single rows on the heatmaps (Figures 1C, 3C, 3H, 4C, 5C, 5H, 5K, S1A, S1C, S1D, S1H, S1K, S1N, S1Q, S4C, S5B). 7) Normalization of each single-nucleus profile by its max intensity value. 8) Averaging and standard error calculation of the single-nuclei profiles within the same experimental conditions to obtain single-condition average profiles and its associated standard errors. Since 3D segmentation is quite time-consuming, we set out to determine if 2D segmentation would yield similar results. Therefore, we segmented the same nuclei in 3D and in 2D, where 3D segmentation allows for automatic determination of the center z-slice (using the z coordinate of the 3D object’s centroid). We then compared the intensity profiles between 3D and 2D segmentation and found minimal differences. Thanks to this validation, we could proceed to segment all the other experiments in 2D.

To obtain the intensity profiles shown in fig. S1S, we applied the profile analysis described above but we included the nucleoli segmentation masks (see section “Nucleolar volume analysis”). The center of the inner dark area was therefore replaced with the centroid of the segmented nucleoli. Finally, the distances were not normalized as in step 4, but they were plotted as absolute differences from the nucleolus edge (in μm).

#### Nucleolar volume analysis

Intestinal nuclei were manually segmented in 2D using Cell-ACDC (*49*) based on HIS-72-GFP or Hoechst 33342 signal and stacked into 3D “cylindrical” objects. To detect and quantify the volume of nucleoli in 3D, we adapted the spot detection routine developed in (*50*) and (*51*). The analysis steps are the following: 1) Application of a 3D gaussian filter with a small sigma (0.75 voxel) of the FIB-1::mCherry signal. 2) Instance segmentation of the spots’ signal using the best-suited automatic thresholding algorithm (either the threshold yen or Li algorithms from the Python library scikit-image (*52*)). 3) 3D connected component labelling of the thresholded mask obtained in step 3 to separate and label the objects. 4) Manual inspection and removal of nucleoli segmented from other tissues. This step is required because nuclei were segmented in 2D and stacked into 3D, raising the possibility of nucleoli from other tissues such as hypoderm to be detected. 5) Volume calculation as the sum of all the voxels belonging to the segmented nucleoli.

#### Positioning of heterochromatic and euchromatic reporters

Quantitation of heterochromatic (fig. S2E) and euchromatic reporter (Fig. 2B) distributions on focal stacks of images was done with the ImageJ plugin PointPicker (http://bigwww.epfl.ch/thevenaz/pointpicker/) as previously described (*13*). Briefly, for fig. S2E, the disc of the nucleus in which the spot is the brightest is divided into three zones of equal surface, each containing 33% of the area and the frequency of the spot in these three zones is quantified for fed and fasted nuclei. For Fig. 2B, the relative nucleolus edge-nuclear periphery distance is divided into 10 bins, with 0-1 closest to the nucleolus edge and 9-10 closest to the nuclear periphery and the frequency of the spot in each bin is scored for fed and fasted nuclei.

### Heat and cold stress

Fed L1 larvae were placed onto standard NGM plates with OP50 and were subjected to either a 6-hour heat shock at 34 °C or a 6-hour cold shock at 6 °C in an air incubator.

### Osmolarity

The final osmolarity of a solution was calculated by summing the osmolarity contributed by each solute present in the solution. To determine the osmolarity of each solute, the molarity of the solute was multiplied by the number of osmoles it produces when dissolved in water. M9 buffer was calculated to be 360 mOsm. To achieve different osmolarities, M9 buffer was diluted in milli Q water to obtain buffers with osmolarities of 180, 150, and 120 mOsm, respectively. To create a solution of 540 mOsm, NaCl (90 mM) was added to M9.

### Actinomycin D treatment

A stock solution of Actinomycin D (Bioaustralis) was prepared at a concentration of 25 mg/ml in DMSO. For the treatment plates, Actinomycin D was diluted into NGM medium to achieve a final concentration of 100 μg/ml. Control plates were prepared with an equivalent amount of DMSO (0.4%). The plates were seeded with OP50 bacteria, and the bacteria were subsequently stained with Hoechst 33342 dye, as previously described. Fed L1s were placed onto Actinomycin D or DMSO plates and imaged after 6 hours.

### Statistics

All statistical tests, exact p values, n and number of biological replicas are listed in table S2.

---

**Fig. S1.**
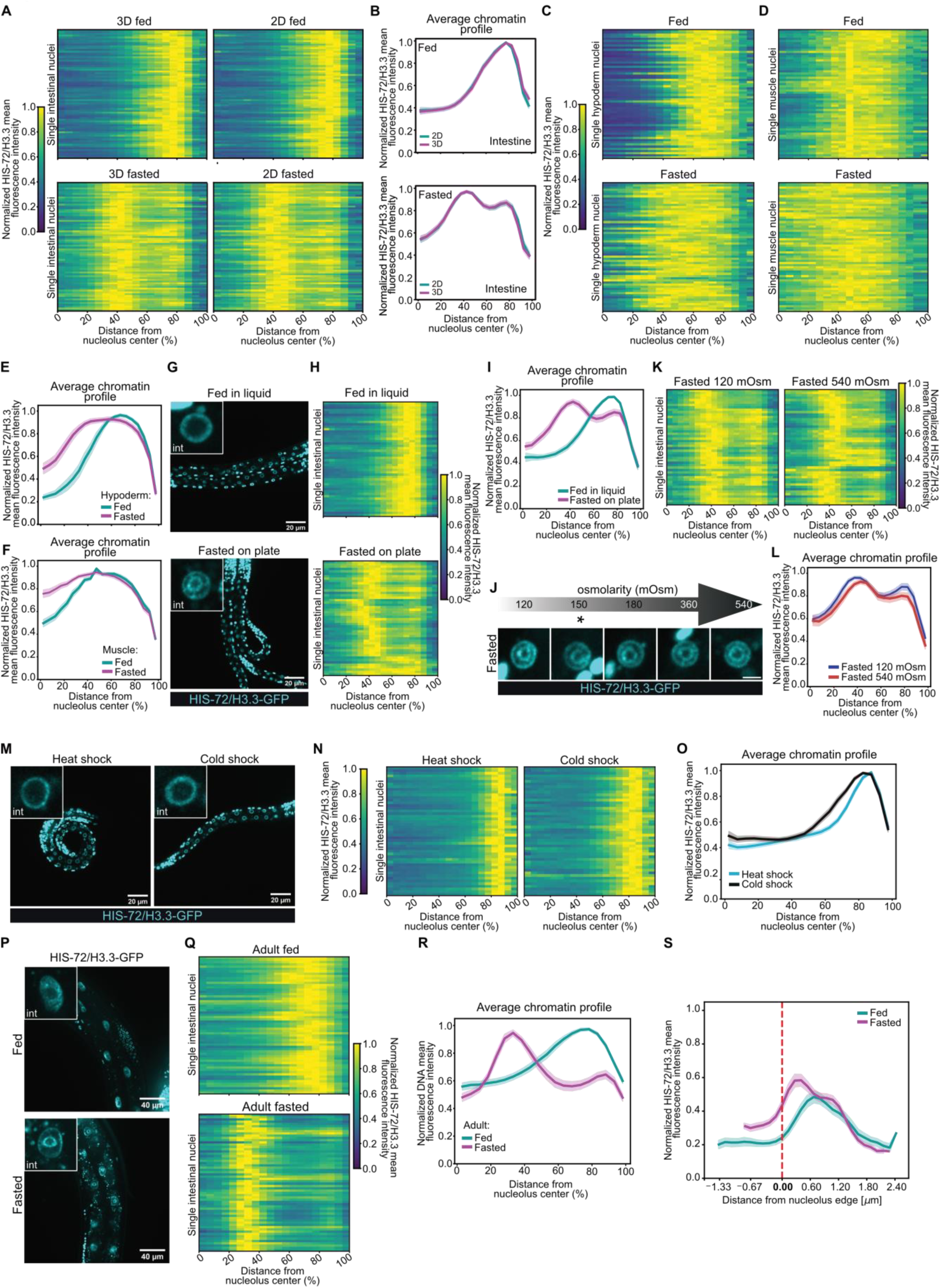
***Related to*** Fig. 1 (**A**) Heatmaps showing the radial fluorescence intensity profiles of HIS-72/H3.3-GFP in intestinal nuclei of wt L1s of the indicated nutritional status as a function of the relative distance from the nucleolus center. Each row corresponds to a single nucleus, segmented in 3D (left) or in 2D at its central plane (right). Radial fluorescence intensity profiles were averaged over all angles. The same 72 intestinal nuclei in animals from 3 independent biological replicas were analyzed for the 3D and 2D segmentation. (**B**) Line plots of the averaged single nuclei profile shown in (A). The shaded area represents the 95% confidence interval of the mean profile. (**C**) Heatmaps as in (A, right) but for hypoderm nuclei. 60 intestinal nuclei were analyzed in animals from 3 independent biological replicas. (**D**) Heatmaps as in (A, right) but for muscle nuclei. 60 intestinal nuclei were analyzed in animals from 3 independent biological replicas. (**E**) Line plots as in (B) but of the averaged single nuclei profile shown in (C). (**F**) Line plots as in (B) but of the averaged single nuclei profile shown in (D). (**G**) Single focal planes of representative wt L1 larvae expressing HIS-72/H3.3-GFP, fed in liquid (top) or fasted on plate (bottom). Insets: zoom of single intestinal nucleus. (**H**) Heatmaps as in (A, right) but for single intestinal nuclei of wt animals fed in liquid (top) or fasted on plate (bottom). 48 intestinal nuclei were analyzed in animals from 3 independent biological replicas. (**I**) Line plots as in (B) but of the averaged single nuclei profile shown in (H). (**J**) Single focal planes of representative, wt intestinal nuclei expressing HIS-72/H3.3-GFP in L1 larvae that were fasted at the indicated osmolarity. * indicated the standard osmolarity of worm plates (150 mOsm). Scale bar indicates 2.5 µm. (**K**) Heatmaps as in (A, right) but for single intestinal nuclei of wt animals fasted at 120 mOsm (left) or at 540 mOsm (right). 48 intestinal nuclei were analyzed in animals from 3 independent biological replicas. (**L**) Line plots as in (B) but of the averaged single nuclei profile shown in (K). (**M**) Single focal planes of representative fed wt L1 larvae expressing HIS-72/H3.3-GFP, heat shocked at 34 °C (left) or cold shocked at 6 °C (right) for 6 hours. Insets: zoom of single intestinal nucleus. (**N**) Heatmaps as in (A, right) but for single intestinal nuclei of wt L1 larvae heat shocked (left) or cold shocked (right). 48 intestinal nuclei were analyzed in animals from 2 independent biological replicas. (**O**) Line plots as in (B) but of the averaged single nuclei profile shown in (N). (**P**) Single focal planes of representative wt adults expressing HIS-72/H3.3-GFP, fed (top) or fasted for 12 hours (bottom). Insets: zoom of single intestinal nucleus. (**Q**) Heatmaps as in (A, right) but for single intestinal nuclei of wt adults fed (top) or fasted (bottom). 60 intestinal nuclei were analyzed in animals from 3 independent biological replicas. (**R**) Line plots as in (B) but of the averaged single nuclei profile shown in (Q). (**S**) Line plots of the averaged distance of the HIS-72/H3.3-GFP signal from the nucleolus edge, identified using FIB-1-mCherry, in single intestinal nuclei of fed and fasted L1 larvae. The shaded area represents the 95% confidence interval of the mean profile. 72 intestinal nuclei were analyzed in animals from 3 independent biological replicas.

**Fig. S2.**
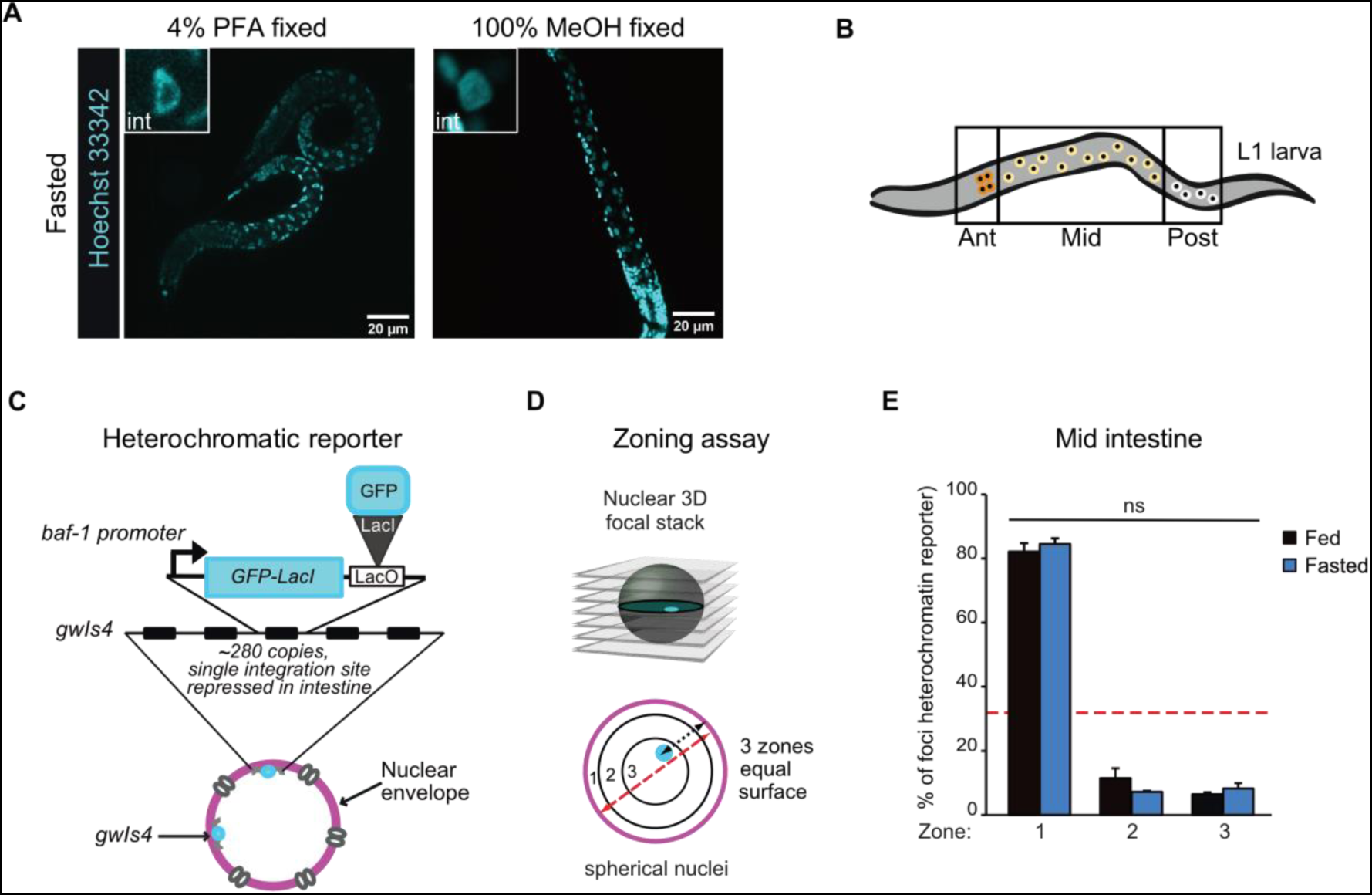
***Related to*** Fig. 2 (**A**) Single focal planes of representative wt L1 larvae fasted, fixed with the indicated chemical and stained with Hoechst 33342 to visualize DNA. (**B**) Schematic representation of intestinal nuclei in L1 larvae. (**C**) Schematic representation of the heterochromatic reporter used in (E). (**D**) Zoning assay for GFP-LacI marked heterochromatic reporter distribution. Radial position is determined relative to the fluorescently-tagged nuclear membrane, and values are binned into three concentric zones of equal surface. Zone 1 is the most peripheral. (**E**) Heterochromatic reporter distribution quantitation, as described in (D), in mid intestine cells of wt fed and fasted L1 larvae. Red dashed line indicates random distribution of 33%. Animals from 3 independent biological replicas, with at least 70 GFP-LacI spots per replica, were analyzed. Error bars are SEM. By χ2 test, fed and fasted samples are not significantly (ns) different from each other. p value and exact n are in table S2.

**Fig. S3.**
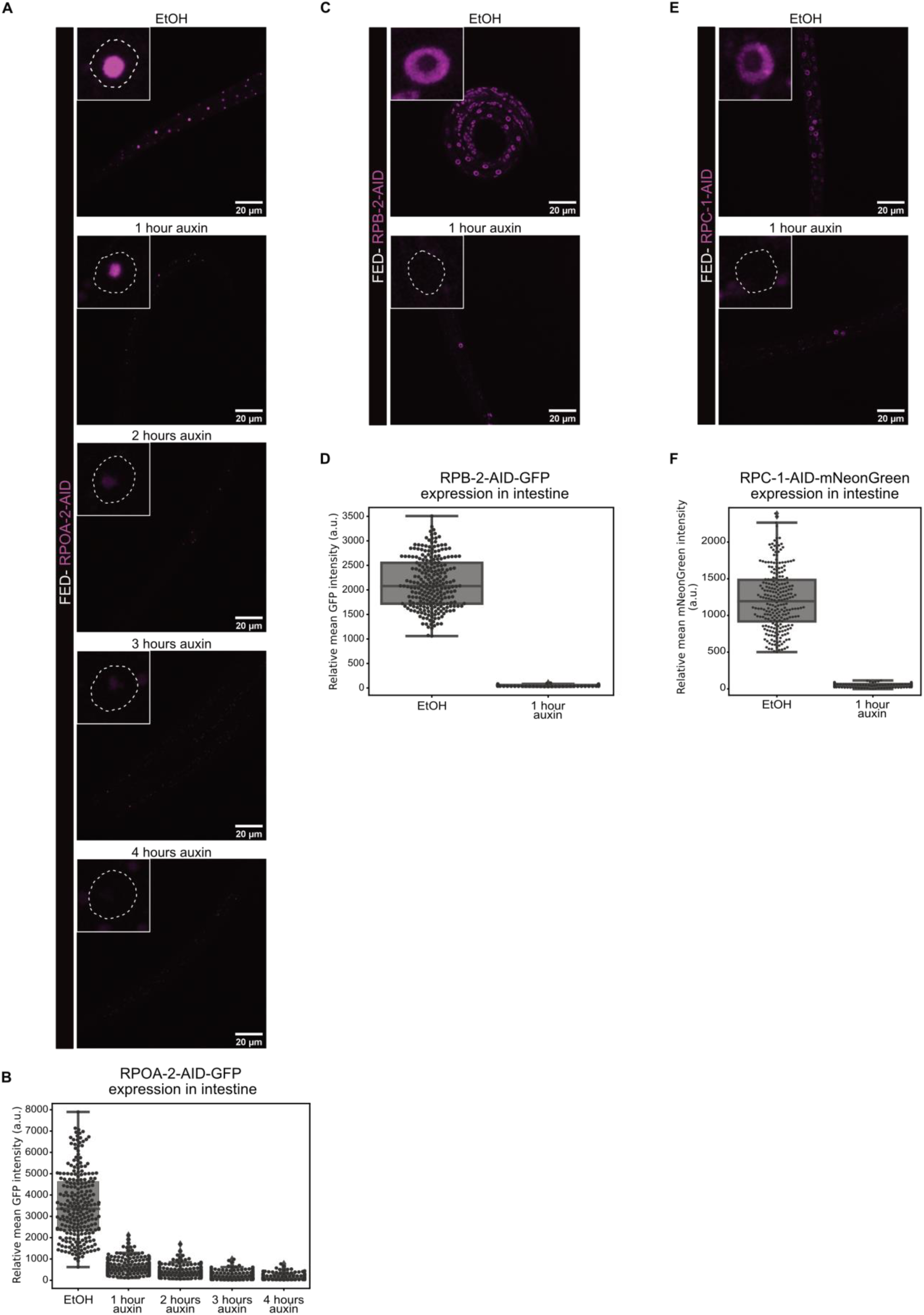
***Related to*** Figs. 3 and 4 (**A**) Single focal planes of representative fed L1 larvae expressing RPOA-2-AID-GFP from its endogenous locus, and TIR1, control treated (EtOH), or treated with 1 mM auxin for the indicated amount of time. Insets: zoom of single intestinal nucleus. (**B**) Boxplots comparing the mean fluorescence intensity of RPOA-2-AID-GFP in intestinal nuclei in fed L1s expressing RPOA-2-AID-GFP and TIR1, control treated (EtOH), or treated with 1 mM auxin for the indicated amount of time. 240 intestinal nuclei were analyzed per condition, from 2 independent biological replicas. (**C**) As in (A), but for animals expressing RPB-2-AID-GFP. (**D**) As in (B), but for animals expressing RPB-2-AID-GFP. (**E**) As in (A), but for animals expressing RPC-1-AID-mNeonGreen. (**F**) As in (B), but for animals expressing RPC-1-AID-mNeonGreen.

**Fig. S4.**
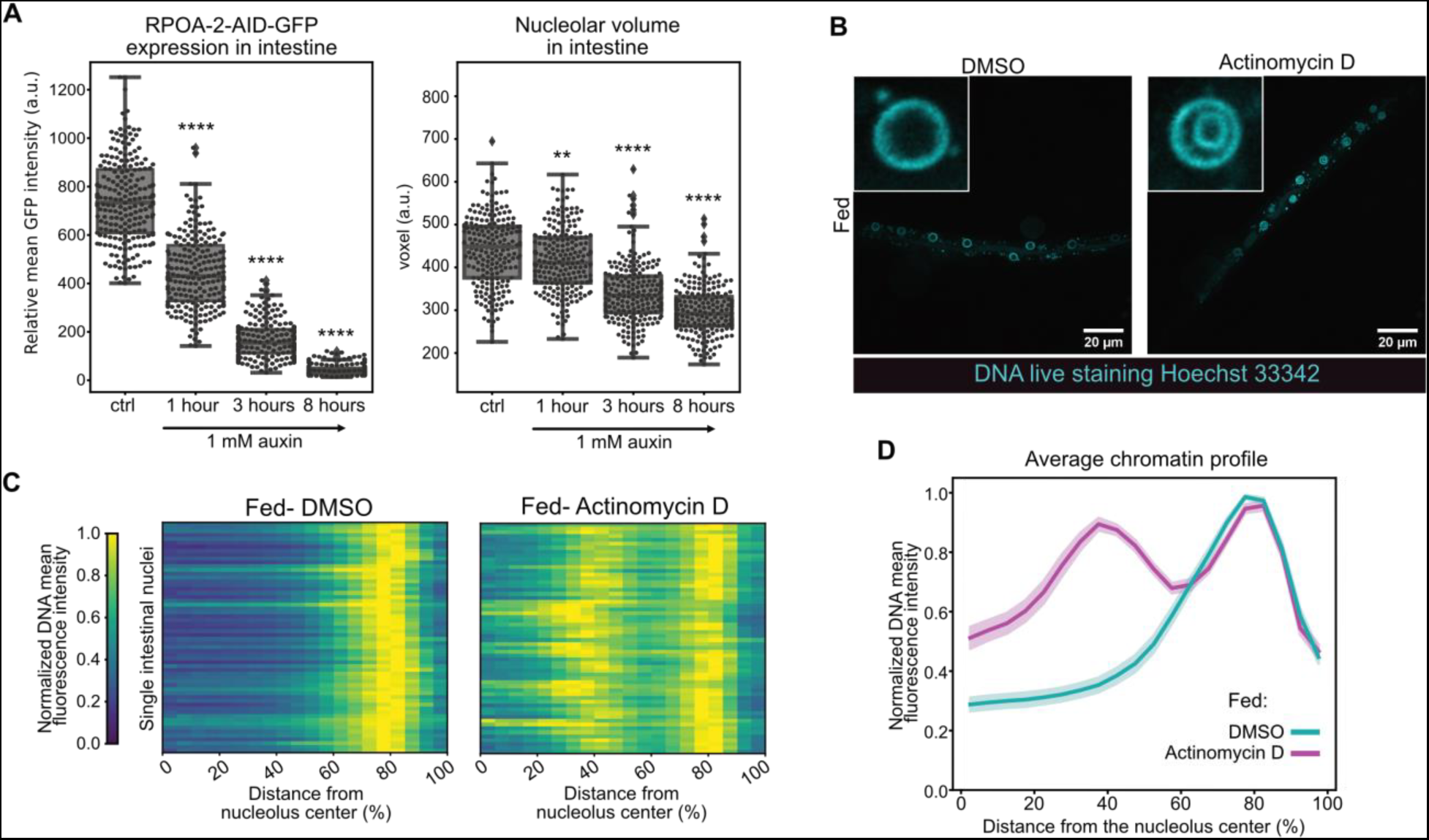
***Related to*** Fig. 4 (**A**) Boxplots comparing the expression of RPOA-2-AID-GFP (left) and volume of the nucleolus, measured with FIB-1-mCherry, (right) in the intestine of fed L1 larvae expressing TIR1-BFP and treated with 1 mM auxin for the indicated time. Probability values from Wilcoxon rank sum tests comparing to before auxin: **** and ** indicate p value <0,0001 and <0,01, respectively. See p values in table S2. **(B**) Single focal planes of representative fed wt L1 larvae, live-stained with Hoechst 33342, treated with DMSO, as control, (left) or Actinomycin D (right). Insets: zoom of single intestine nucleus. (**C**) Heatmaps showing the radial fluorescence intensity profiles of Hoechst 33342-stained DNA in intestinal nuclei of fed wt animals treated with DMSO, as control (left), or Actinomycin D (right), as a function of the relative distance from the nucleolus center. Each row corresponds to a single nucleus, segmented in 2D at its central plane. Radial fluorescence intensity profiles were averaged over all angles. 60 intestinal nuclei were analyzed in animals from 3 independent biological replicas. (**D**) Line plots of the averaged single nuclei profile shown in (C). The shaded area represents the 95% confidence interval of the mean profile.

**Fig. S5.**
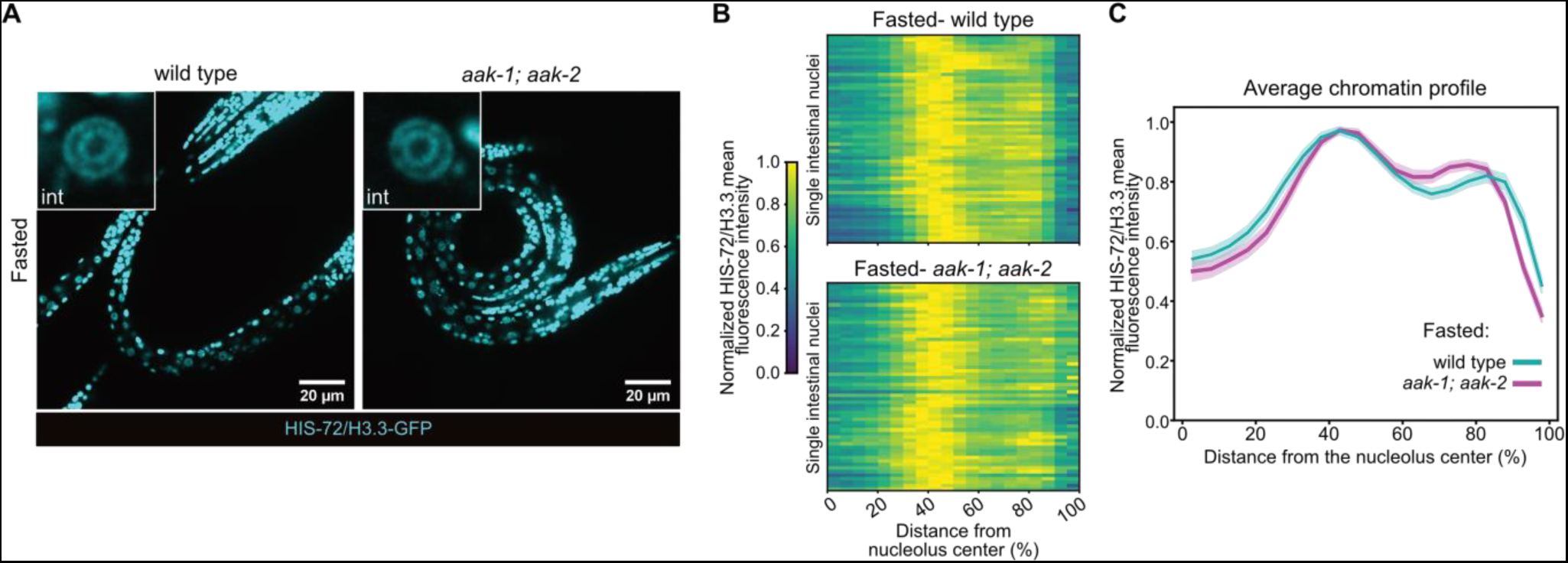
***Related to*** Fig. 5 (A) Single focal planes of representative fasted L1 larvae wild type or double *aak-1* and *aak-2* mutants, expressing fluorescently tagged HIS-72/H3.3. Insets: zoom of single intestine nucleus. (B) Heatmaps showing the radial fluorescence intensity profiles of HIS-72/H3.3-GFP in intestinal nuclei of fasted wild type (top) or *aak-1; aak-2* mutants (bottom) animals, as a function of the relative distance from the nucleolus center. Each row corresponds to a single nucleus, segmented in 2D at its central plane. Radial fluorescence intensity profiles were averaged over all angles. 60 intestinal nuclei were analyzed in animals from 4 independent biological replicas. (C) Line plots of the averaged single nuclei profile shown in (B). The shaded area represents the 95% confidence interval of the mean profile.

**Table S1.**
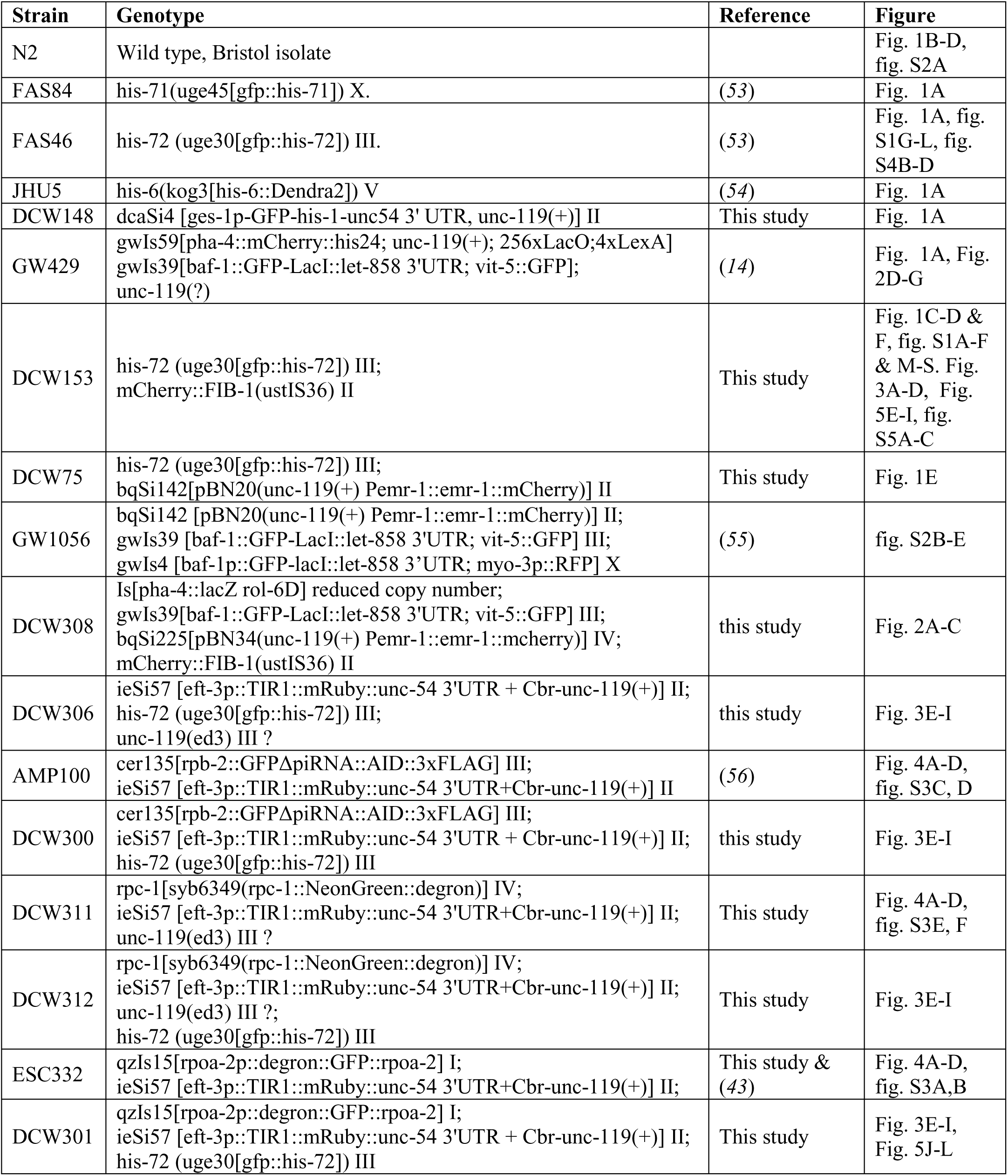
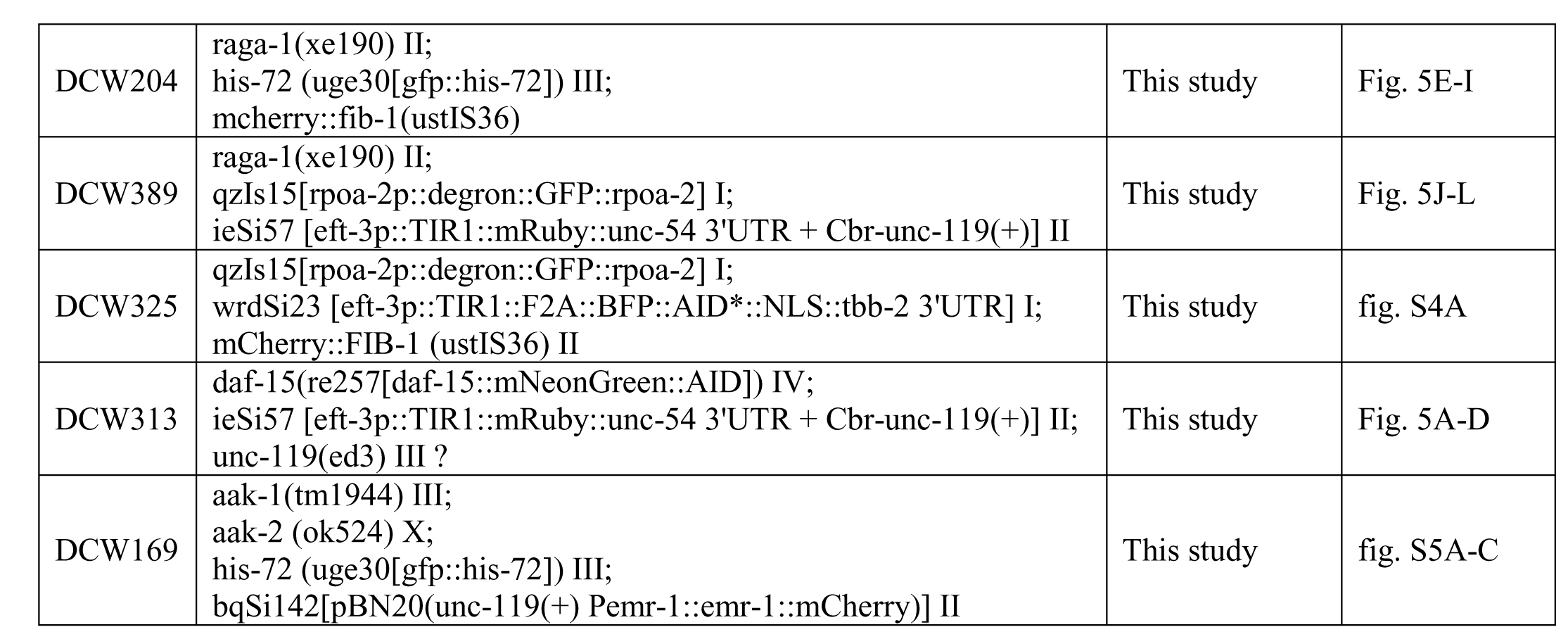
List of worm strains used in this study

**Table S2.**
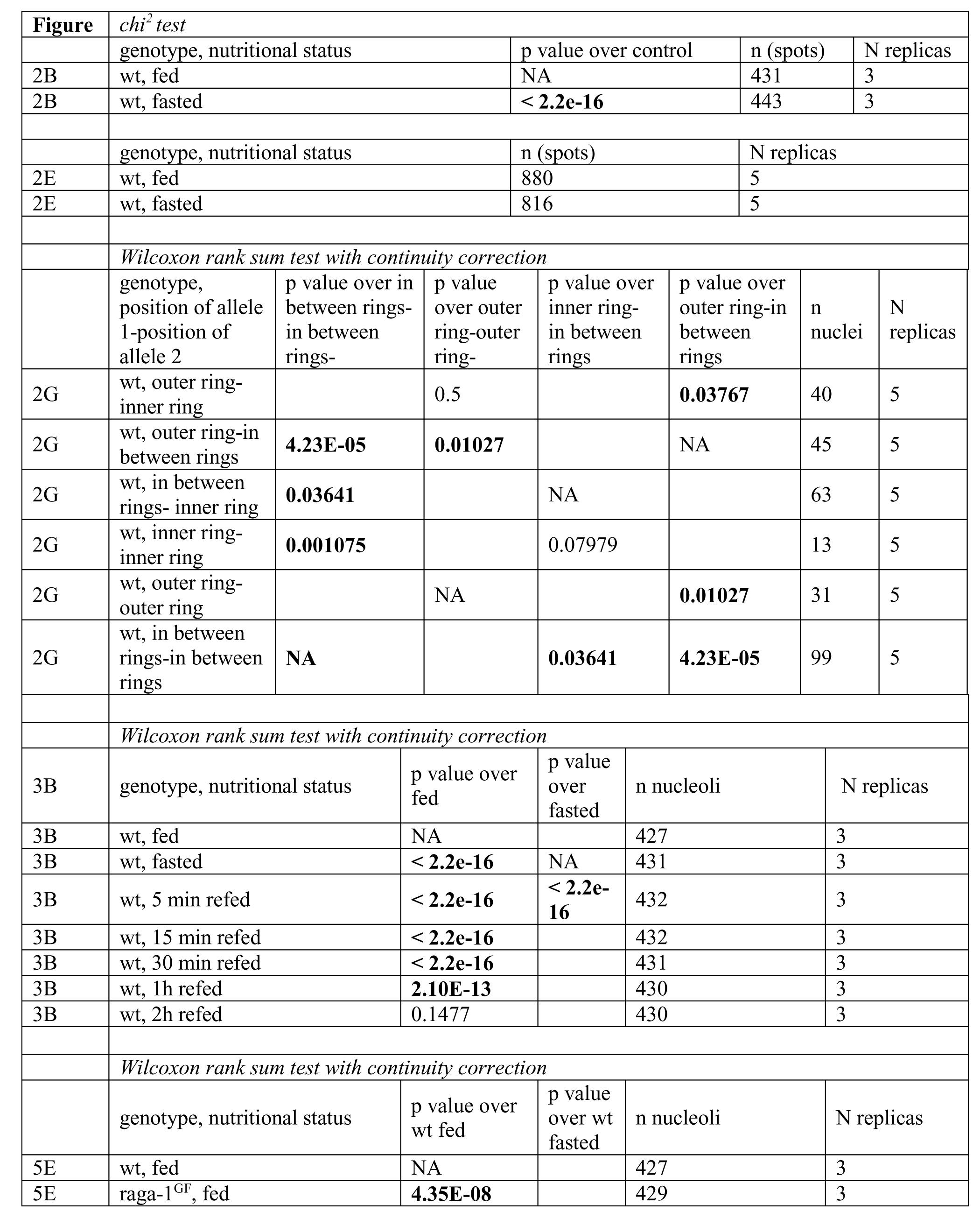
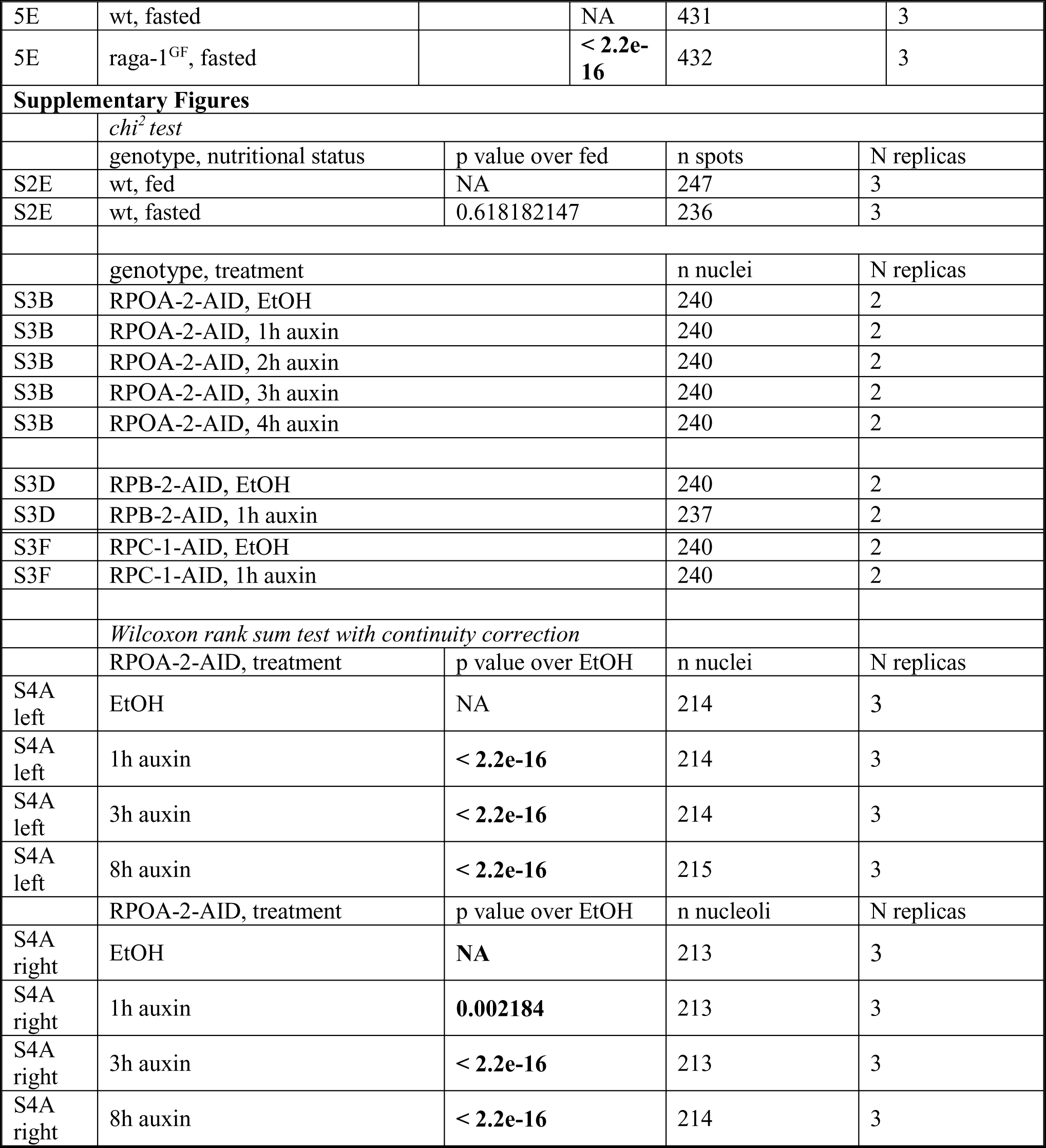
N, p values and number of biological replicas (N) of the experiments in this study.

| Figure | chi<sup>2</sup> test |  |  |  |  |  |  |
| --- | --- | --- | --- | --- | --- | --- | --- |
|  | genotype, nutritional status |  |  | p value over control |  | n (spots) | N replicas |
| 2B | wt, fed |  |  | NA |  | 431 | 3 |
| 2B | wt, fasted |  |  | < **2.2e-16** |  | 443 | 3 |
|  | genotype, nutritional status |  |  | n (spots) |  | N replicas |  |
| 2E | wt, fed |  |  | 880 |  | 5 |  |
| 2E | wt, fasted |  |  | 816 |  | 5 |  |
|  | Wilcoxon rank sum test with continuity correction |  |  |  |  |  |  |
|  | genotype,<br>position of allele<br>1-position of<br>allele 2 | p value over in<br>between rings-<br>in between<br>rings- | p value<br>over outer<br>ring-outer<br>ring- | p value over<br>inner ring-in between<br>rings | p value over<br>outer ring-in<br>between<br>rings | n<br>nuclei | N<br>replicas |
| 2G | wt, outer ring-<br>inner ring |  | 0.5 |  | **0.03767** | 40 | 5 |
| 2G | wt, outer ring-in<br>between rings | **4.23E-05** | **0.01027** |  | NA | 45 | 5 |
| 2G | wt, in between<br>rings- inner ring | **0.03641** |  | NA |  | 63 | 5 |
| 2G | wt, inner ring-<br>inner ring | **0.001075** |  | 0.07979 |  | 13 | 5 |
| 2G | wt, outer ring-outer ring |  | NA |  | **0.01027** | 31 | 5 |
| 2G | wt, in between<br>rings-in between<br>rings | **NA** |  | **0.03641** | **4.23E-05** | 99 | 5 |
|  | Wilcoxon rank sum test with continuity correction |  |  |  |  |  |  |
| 3B | genotype, nutritional status |  | p value over<br>fed | p value<br>over<br>fasted | n nucleoli |  | N replicas |
| 3B | wt, fed |  | NA |  | 427 |  | 3 |
| 3B | wt, fasted |  | < **2.2e-16** | NA | 431 |  | 3 |
| 3B | wt, 5 min refed |  | < **2.2e-16** | < **2.2e-16** | 432 |  | 3 |
| 3B | wt, 15 min refed |  | < **2.2e-16** |  | 432 |  | 3 |
| 3B | wt, 30 min refed |  | < **2.2e-16** |  | 431 |  | 3 |
| 3B | wt, 1h refed |  | **2.10E-13** |  | 430 |  | 3 |
| 3B | wt, 2h refed |  | 0.1477 |  | 430 |  | 3 |
|  | Wilcoxon rank sum test with continuity correction |  |  |  |  |  |  |
|  | genotype, nutritional status |  | p value over<br>wt fed | p value<br>over wt<br>fasted | n nucleoli |  | N replicas |
| 5E | wt, fed |  | NA |  | 427 |  | 3 |
| 5E | raga-1<sup>GF</sup>, fed |  | **4.35E-08** |  | 429 |  | 3 |

|  |  |  |  |  |  |
| --- | --- | --- | --- | --- | --- |
| 5E | wt, fasted |  | NA | 431 | 3 |
| 5E | raga-1<sup>GF</sup>, fasted |  | < **2.2e-16** | 432 | 3 |
| Supplementary Figures |  |  |  |  |  |
|  | *chi<sup>2</sup> test* |  |  |  |  |
|  | genotype, nutritional status | p value over fed |  | n spots | N replicas |
| S2E | wt, fed | NA |  | 247 | 3 |
| S2E | wt, fasted | 0.618182147 |  | 236 | 3 |
|  | genotype, treatment |  |  | n nuclei | N replicas |
| S3B | RPOA-2-AID, EtOH |  |  | 240 | 2 |
| S3B | RPOA-2-AID, 1h auxin |  |  | 240 | 2 |
| S3B | RPOA-2-AID, 2h auxin |  |  | 240 | 2 |
| S3B | RPOA-2-AID, 3h auxin |  |  | 240 | 2 |
| S3B | RPOA-2-AID, 4h auxin |  |  | 240 | 2 |
| S3D | RPB-2-AID, EtOH |  |  | 240 | 2 |
| S3D | RPB-2-AID, 1h auxin |  |  | 237 | 2 |
| S3F | RPC-1-AID, EtOH |  |  | 240 | 2 |
| S3F | RPC-1-AID, 1h auxin |  |  | 240 | 2 |
|  | *Wilcoxon rank sum test with continuity correction* |  |  |  |  |
|  | RPOA-2-AID, treatment | p value over EtOH |  | n nuclei | N replicas |
| S4A left | EtOH | NA |  | 214 | 3 |
| S4A left | 1h auxin | < **2.2e-16** |  | 214 | 3 |
| S4A left | 3h auxin | < **2.2e-16** |  | 214 | 3 |
| S4A left | 8h auxin | < **2.2e-16** |  | 215 | 3 |
|  | RPOA-2-AID, treatment | p value over EtOH |  | n nucleoli | N replicas |
| S4A right | EtOH | NA |  | 213 | 3 |
| S4A right | 1h auxin | **0.002184** |  | 213 | 3 |
| S4A right | 3h auxin | < **2.2e-16** |  | 213 | 3 |
| S4A right | 8h auxin | < **2.2e-16** |  | 214 | 3 |

